# The Spectrum of Asynchronous Dynamics in Spiking Networks: A Theory for the Diversity of Non-Rhythmic Waking states in Neocortex

**DOI:** 10.1101/427765

**Authors:** Yann Zerlaut, Stefano Zucca, Stefano Panzeri, Tommaso Fellin

## Abstract

The cerebral cortex of awake animals exhibits frequent transitions between diverse non-rhythmic network states. However, it is still unclear how these different activity states emerge within the same network and how each state impacts network function. Here, we demonstrate that model networks of spiking neurons with moderate recurrent interactions dynamically change their asynchronous dynamics depending upon the level of afferent excitation. We found that the model network displayed a spectrum of asynchronous states, ranging from afferent input-dominated (AD) regimes, characterized by unbalanced synaptic currents and sparse firing, to recurrent input-dominated (RD) regimes, characterized by balanced synaptic currents and dense firing. The model predicted regime-specific relationships between several different neural biophysical properties which were all experimentally confirmed by intracellular recordings in the somatosensory cortex of awake mice. Moreover, theoretical analysis showed that AD regimes more precisely encode spatiotemporal patterns of presynaptic activity, while RD regimes better encoded the strength of afferent inputs. These results provide a theoretical foundation for how recurrent neocortical circuits generate non-rhythmic waking states and how these different states modulate the processing of incoming information.

## Introduction

Cortical circuits frequently display spontaneous activities which are characterized by small values of pairwise spiking synchrony (Ecker et al., 2010; Renart et al., 2010) and which are generally referred to as asynchronous dynamics. The current theoretical description of such asynchronous regimes is based on an emergent solution of balanced synaptic activity in recurrently connected networks (Amit and Brunel, 1997; Destexhe and Contreras, 2006; Kumar et al., 2008; Litwin-Kumar and Doiron, 2012; Parga, 2013; Renart et al., 2010; Tsodyks and Sejnowski, 1995; Vogels et al., 2005; van Vreeswijk and Sompolinsky, 1996). In this dynamical setting, the strong excitatory and inhibitory currents cancel each other to bring the average membrane potential (V_m_) below the spiking threshold, and the intense synaptic bombardment results in Gaussian V_m_ fluctuations (van Vreeswijk and Sompolinsky, 1996). Early recordings in awake cats supported this picture: V_m_ dynamics during desynchronized activity displayed near-Gaussian fluctuations and tonic membrane depolarization bringing the neuron close to the spiking threshold (Steriade et al., 2001). However, recent experiments in awake behaving rodents suggested a more complex picture (Busse et al., 2017; McGinley et al., 2015a; Nakajima and Halassa, 2017). Together with rhythmic activity in the [2,10] Hz range (Crochet and Petersen, 2006; Poulet and Petersen, 2008), spontaneous dynamics exhibited diverse asynchronous states (non-rhythmic activity) characterized by different mean V_m_ (McGinley et al., 2015b; Polack et al., 2013; Reimer et al., 2014) and firing activity (Vinck et al., 2015).

At the theoretical level, those observations raise fundamental questions. What is the dynamical nature of the different asynchronous states of wakefulness? Is the classical setting of a recurrently-balanced dynamics a valid description for all the different asynchronous states? If not, does asynchronous dynamics exist beyond such a setting? How can we develop a tractable computational model that reveals the mechanisms generating these asynchronous states, that explains quantitively the membrane potential dynamics observed in cortex during wakefulness, and that allows us to understand the specific computational advantages of each state?

To address the above questions, we explored the collective dynamics emerging in recurrently connected networks of excitatory and inhibitory spiking units. We found that, for moderate recurrent interactions, spiking network models displayed a spectrum of stable asynchronous states exhibiting spiking activity spanning over orders of magnitudes, and we found profoundly different contributions of the afferent and recurrent components in shaping network dynamics. The model made precise predictions for a number of relationships among different electrophysiologically-measurable features which were all experimentally confirmed by electrophysiological recordings of neural activity across different non-rhythmic states of waking in the superficial layers of the mouse primary somatosensory cortex. Moreover, we demonstrated, in network models, that different activity regimes were characterized by distinct stimulus information coding properties. These results provide a new theoretical framework for explaining the origin and the information coding properties of the diverse non-rhythmic states observed during wakefulness.

## Results

### Recurrent networks exhibit a spectrum of asynchronous regimes upon modulation of their level of afferent excitation

We hypothesized that the various non-rhythmic regimes of wakefulness could be described by a set of emergent solutions of recurrent activity in excitatory/inhibitory spiking networks. More specifically, we reasoned that regimes of intense synaptic bombardments (Brunel, 2000; Kumar et al., 2008; Renart et al., 2010; van Vreeswijk and Sompolinsky, 1996) should be complemented with regimes of low spiking activity, where single neurons dynamics is driven by a few synaptic events, to describe the lower depolarization which characterizes asynchronous regimes associated to moderate arousal (McGinley et al., 2015a). Consequently, to find such a set of solutions, we explored the dynamics of spiking networks in a wide activity range including low levels of recurrent activity (< 0.1 Hz). We implemented this in a randomly connected recurrent network of leaky integrate-and-fire excitatory and inhibitory neurons with conductance-based synapses (Kumar et al., 2008). The model network incorporated the following experimentally-driven features: recurrent synaptic weights leading to post-synaptic deflections below 2 mV at rest (Jiang et al., 2015; Lefort et al., 2009; Markram et al., 2015); probabilities of connectivity among neurons matching the relatively sparse ones observed in the adult mouse sensory cortex (Jiang et al., 2015) (see also **Discussion**); a potent external afferent input to account for the numerous and synchronized excitatory thalamic drives onto sensory cortices (Bruno and Sakmann, 2006); and a higher excitability of inhibitory cells to model the high firing of a subpopulation of interneurons, the fast-spiking non-adapting neurons (Markram et al., 2004). The network model and its features are depicted in **Figure 1A**. All parameters are listed in **Table 1**.

**Figure 1.**
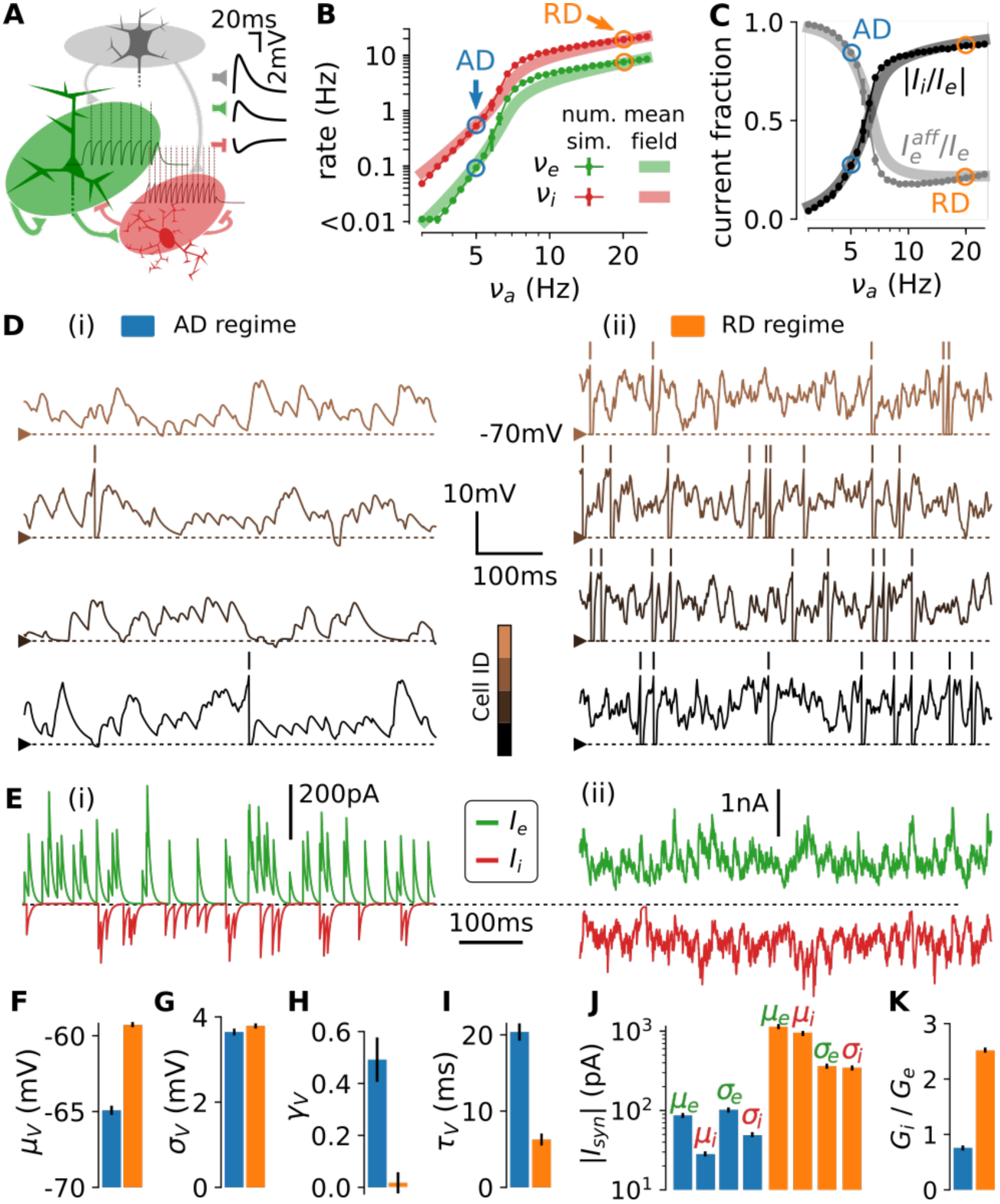
A spectrum of asynchronous regimes emerges in a recurrent spiking network upon modulation of its level of afferent excitation. **(A)** The reduced model of the neocortical assembly. An afferent excitatory input targets the recurrently connected excitatory (green) and inhibitory (red) populations. In the inset, we show the post-synaptic deflections at rest (−70 mV) associated to each type of synaptic connection (grey indicates the afferent population). We also show the spiking response of single neurons to a current pulse of 120 pA (background traces). Note the higher excitability of inhibitory cells because of their lower AP threshold (−53 mV *vs* −50 mV for excitatory cells). Parameters of the model are listed in **Table 1**. **(B)** Stationary firing rates of the excitatory (green) and inhibitory (red) populations as a function of the level of afferent activity (dots with error bars represent mean±s.e.m over n = 10 numerical simulations). The AD (blue circle) and the RD (orange circle) levels are indicated. We show the mean-field predictions for comparison (thick transparent lines, see main text and **Figures S2**). **(C)** Fraction of afferent currents within the sum of recurrent and afferent excitatory currents 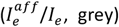 and absolute ratio between inhibitory and excitatory currents (|*I_i_*/*I_e_*|, black) as a function of the level of afferent activity *v_a_* (dots with error bars: n = 10 simulations; thick transparent lines: mean-field predictions). **(D)** Membrane potential traces for four neurons in the AD (i) and in the RD (ii) regimes. **(E)** Excitatory (green) and inhibitory (red) synaptic currents targeting a single neuron in the AD (i, left) and RD (ii, right) regimes. Note the difference in current-scale between (i) and (ii). **(F-K)** Electrophysiological signature of the two regimes (blue and orange for AD and RD respectively) at the single cell level (evaluated on excitatory cells on a single simulation, error bars represent variability over n = 10 recorded cells). (**F**) mean depolarization *μ_V_*, (**G**) standard deviation *σ_V_*, (**H**) skewness of the *V_m_* distribution *γ_V_*, (**I**) autocorrelation time *τ_V_*, (**J**) mean and standard deviation (*μ* and *σ* respectively) of the excitatory (indexed by *e*) and inhibitory currents (indexed by *i*) over time, and (**K**) ratio of inhibitory to excitatory synaptic conductances *G_i_*/ *G_e_*.

**Table 1.**
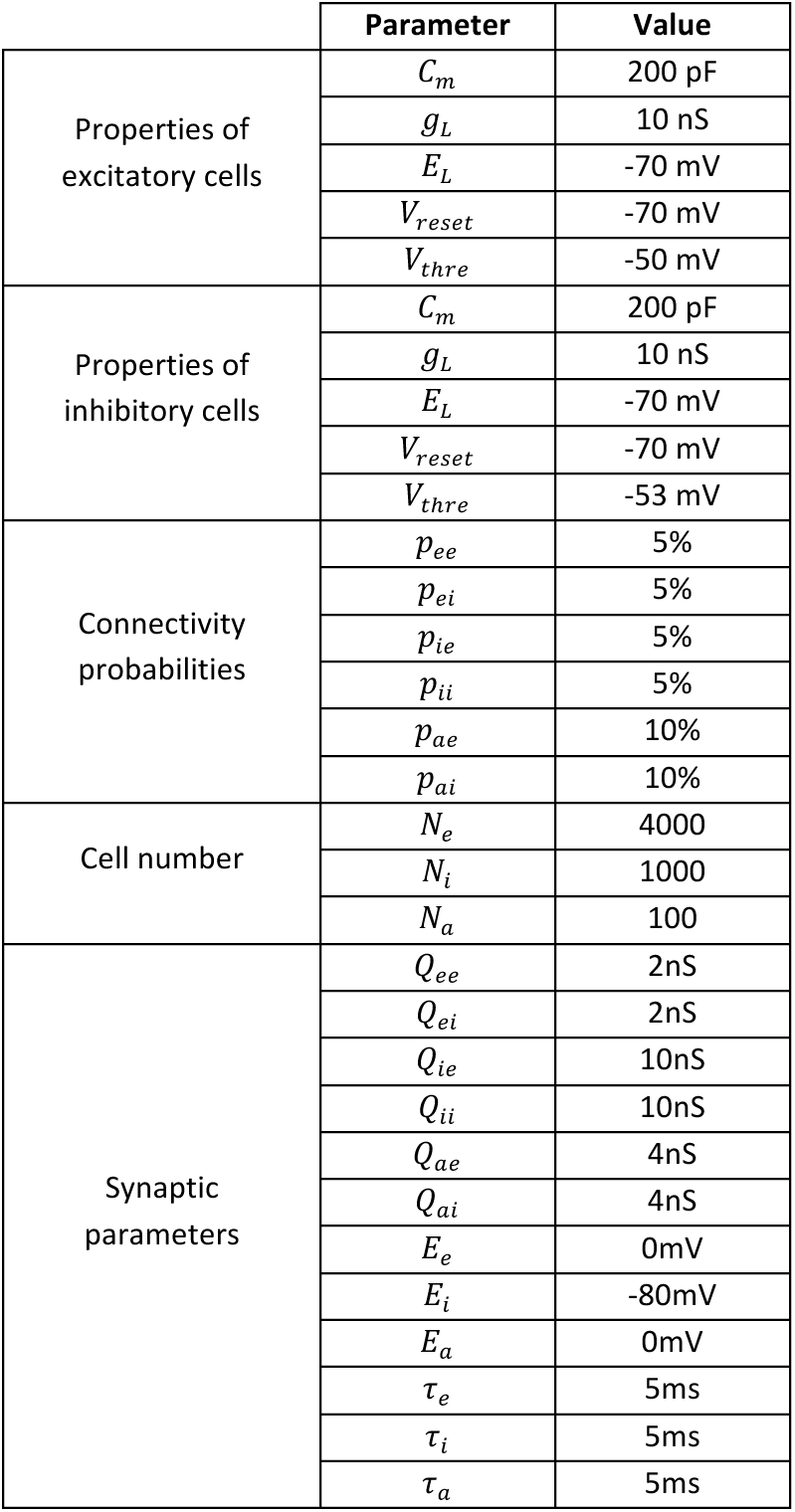
Network parameters for the two population
model (excitation/inhibition) with afferent excitation.

We tested the above hypothesis by analyzing the emergent network dynamics as a function of the stationary level of afferent excitation. We found that the model produced stable asynchronous dynamics over a wide range of excitatory and inhibitory activity. The stationary spiking activity of the network spanned four orders of magnitude (**Figure 1B**), while pairwise synchrony remained one order of magnitude below classical synchronous regimes (*SI* < 5e-3, see **Figure S1** for a detailed analysis of the network’s residual synchrony). Varying the model’s afferent population activity, *v_a_*, from *v_a_* = 3 Hz to *v_a_* = 25 Hz resulted in a logarithmically-graded increase of excitatory firing rates, *v_e_*, from *v_e_* = 0.004 Hz to *v_e_* = 8.5 Hz and inhibitory firing rates, *v_i_*, from *v_i_* = 0.07 Hz to *v_i_*)= 21.8 Hz (**Figure 1B**). Thus, recurrent dynamics performed an exponentiation of the level of afferent input. Importantly, the relative contributions of the afferent and recurrent excitation in shaping the single neuron dynamics varied over the different levels of activity (grey curve in **Figure 1C**), it varied from regimes dominated by the afferent excitation 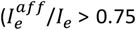 for *v_a_* ≤ 6 Hz, where *I_e_* (is the sum of the afferent 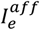 and recurrent 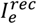 excitatory currents) to a recurrent connectivity-dominated regime 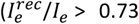 *v_a_* ≥ 12 Hz). Concomitantly, the ratio between mean inhibitory and excitatory synaptic currents (*I_i_*/*I_e_*, where *I_i_*) is the recurrent inhibitory current) was not constant over those different activity levels (black curve in **Figure 1C**). Rather, it gradually varied from excitatory-dominated regimes where *I_i_*/*I_e_* ≪ 1 *I_i_*/*I_e_* < 0.50 below for *v_a_*= 6 Hz) to balanced activity where *I_i_*/*I_e_* ~ 1(*I_i_*/*I_e_* > 0.85 for *v_a_* ≥ 12 Hz). In the following, we refer to this continuum of diverse emergent solutions of recurrent activity as a “spectrum” of asynchronous regimes.

To better highlight the profound differences in terms of network dynamics and of electrophysiologically measurable neural activity features that are observed along the spectrum, we selected two levels of afferent drive leading to two relative extreme states along this spectrum (**Figure 1B**). The first example state, termed Afferent input-Dominated state and shortened to “AD”, was a state found at low afferent excitation that, as we shall show below, was characterized by temporally sparse spiking activity and was dominated by its afferent excitation. The second example state, termed Recurrent input-Dominated state and shortened to “RD”, was a state found at high afferent excitation that, as we shall show below, was characterized by temporally dense spiking activity and was dominated by its synaptically balanced recurrent activity. We show samples of membrane potential traces (**Figure 1D**) and synaptic currents (**Figure 1E**) for the two selected regimes in representative simulated neurons.

For high afferent excitation (*v_a_* = 20 Hz, RD), recurrent activity was dense (> 1 Hz, here *v_e_* = 7.6 ± 0.1 Hz and *v_i_* = 19.2 ± 0.2 Hz), and the network displayed asynchronous dynamics in the balanced setting. The electrophysiological features identifying this regime were: 1) a mean depolarized *V_m_* (*μ_V_* = −59.3 + 0.1 mV, **Figure 1F**) with a standard deviation (*σ_V_*= 3.7 ± 0.1 mV, **Figure 1G**) which implied that fluctuations in V_m_ bring V_m_ close to the spiking threshold (*V_thre_* = −50 mV) (see *V_m_* traces in **Figure 1D**); 2) a symmetric *V_m_* distribution (see the near-zero skewness in **Figure 1H**, *γ_V_* = 0.02 ± 0.04) that was a signature of Gaussian fluctuations (coefficient of determination of a Gaussian fitting after blanking spikes: *R*^2^ = 0.994±0.002); 3) fast membrane potential fluctuations (the autocorrelation time of the fluctuations *τ_V_* = 6.2 ± 0.8 ms was much lower than the membrane time constant at rest 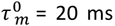, **Figure 1I**); 4) a high conductance state (synaptic conductances sum up to more than four times the leak conductance (Destexhe et al., 2003), here the conductance ratio was 6.6 ± 0.1); 5) balanced excitatory and inhibitory currents (|*I_i_*)/*I_e_*| = 0.881 ± 0.003, **Figure 1C**) with large means compared to their temporal fluctuations (*μ_e_*/*σ_e_* = 3.2 ± 0.1 and *μ_i_*/*σ_i_*) = 2.8 ± 0.1, **Figure 1J**); 6) the predominance of the recurrent activity in shaping single neuron dynamics (recurrently-mediated synaptic currents represented 84.6 ± 0.1% of the membrane currents and largely exceeded the contributions of the afferent excitatory currents and leak currents, 10.8 ± 0.1% and 4.6 ± 0.1% respectively).

At low levels of afferent activity (*v_a_* = 5 Hz, AD), a stable regime of asynchronous dynamics exhibited a qualitatively different set of electrophysiological features. Spiking activity in the network was very sparse (*v_e_* = 0.094 ± 0.008 Hz and *v_i_*= 0.535 ± 0.013 Hz, **Figure 1B**) which, at the single neuron level, was associated to: 1) a higher distance between the mean V_m_ and the spiking threshold (*μ_V_* = 64.1 ± 0.3 mV, **Figure 1F**); 2) a strongly skewed *V_m_* distribution (*γ_V_*: = 0.49 ± 0.09, **Figure 1H**); 3) slower *V_m_* fluctuations (*τ_V_* = 20.4 ± 1.1 ms, i.e. a threefold increase with respect to the RD regime, **Figure 1I**); 4) a lower conductance state preserving the efficacy of synaptically-evoked depolarizations (instead of the strong shunting characterizing high conductance states, here synaptic conductances increased the input conductance by only 18.2 ± 1.0 %); 5) excitatory dominated synaptic currents, the mean of the excitatory currents largely exceeded those of inhibitory currents (|*I_i_*/*I_e_*| = 0.272 ± 0.018, **Figure 1C**), leading to a nearly equal ratio of conductances (*G_i_/G_e_* = 0.8 ± 0.2, instead of *G_i_/G_e_* = 2.5 ± 0.1 for the balanced currents of RD, **Figure 1K**); 6) the predominance of the non-recurrent components in shaping single neuron dynamics (recurrently-mediated synaptic currents only represented 14.6 ± 0.2% of the membrane currents and were largely inferior to the afferent excitatory current and leak current contributions, 44.9 ± 1.3% and 40.4 ± 1.5% respectively). In contrast to the RD state, the stability of the AD regime did not rely on the excitatory/inhibitory balance of synaptic currents (see **Figure 1E**): the very low amount of recurrent inhibitory currents did not cancel the afferent-dominated sum of excitatory currents (see **Figure 1C,E,J**). Instead, leak currents insured stability by contributing significantly to single neuron integration: the temporal dynamics of the membrane potential was dominated by leak-mediated repolarization following sparse synaptic events (see **Figure 1D**) and accordingly, the autocorrelation time *τ_V_* = 20.4 ± 1.1 ms was close to the membrane time constant at rest 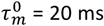. This network state was thus mainly shaped by the following ingredients: a lower frequency excitatory afferent input producing strongly skewed *V_m_* fluctuations (occasionally large enough to elicit spikes) counter-balanced by leak repolarization currents.

### *A mean-field* description including non-Gaussian *V_m_* properties predicts the emergence of the spectrum

The simulations presented above highlight the firing rates (*v_e_, v_i_*) and the *V_m_* fluctuations properties (*μ_V_*, *σ_V_*, *τ_V_*, *γ_V_*,) as key determinants of different states in the spectrum. To understand whether the variations of those statistical quantities provide sufficient ingredients for the emergence of the spectrum, we developed and analyzed a “mean field” description of network activity including these quantities (see **Supplementary Information**). Briefly, a mean field description of network activity reduces the firing rate dynamics of each population into the dynamics of a prototypical neuron whose behavior is captured by a rate-based input-output function; see (Renart et al., 2004) for a review. In standard mean-field approaches, the neuronal input-output function is determined analytically by converting the input firing rates into Gaussian fluctuations of synaptic currents in turn translated into an output firing rate using estimates from stochastic calculus (Tuckwell, 2005). Here, however, building on previous work (Zerlaut et al., 2016), we extended this formalism so that the input firing rates are converted into *V_m_* fluctuations properties that also include higher order non-Gaussian properties (such as the skewness and tail integral of the distribution), and that are in turn converted into an output firing rate thanks to a semi-analytical approach (see **Supplementary Information**). Importantly, we found that the spectrum of activity regimes found in the numerical simulations was also present in such mean-field description (**Figure 1B,C** and **Figure S2).** Because the mean-field description only specifies the firing rates and the four basic properties of *V_m_* statistics that we plotted in our earlier simulations, this analysis further demonstrates that changes in those parameters are sufficient to generate changes in the spectrum. This confirms that the spectrum can be generated also without relying on specific properties in the architecture of the numerical network (such as a degree of clustering within the drawn recurrent connectivity) or more complex dynamical features (such as pairwise synchrony, or deviations from the Poisson spiking statistics), that were not included in the mean field approach. It also strengthen the conclusion that the spectrum dynamics of both real and simulated data can be meaningfully studied and analyzed by quantifying the firing rate and *V_m_* properties.

### A moderate strength of recurrent interactions is a necessary condition for the emergence of the spectrum

Which are the crucial network parameters that lead to the emergence of the spectrum of activity states? We addressed this question through parameter variations in the numerical model.

We first considered what happened when increasing, with respect to the reference network configuration considered above, the value of the excitatory and inhibitory recurrent synaptic weights (**Figure 2**). When multiplying this value by a moderate amount with a factor *f* in the range between 0.1 and 2 (see the depicted examples of *f* = 0.5 and *f* = 1.2, **Figure 2A**), the network was still able to create states of very low (respectively very high) activity at lower (respectively higher) afferent activity. For synaptic weights in this range, the network exponentiated the level of afferent input to generate recurrent activity spanning several orders of magnitude (from 0.004 Hz to 15 Hz, **Figure 2C,D**) with transitions from regimes dominated by afferent inputs 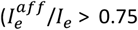 for *v_a_* < 5 Hz, **Figure 2E**) and excitatory currents (|*I_i_*/*I_e_*| < 0.25 for *v_a_* < 5 Hz, **Figure 2F**) to regimes dominated by recurrent activity 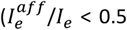 for *v_a_* > 20 Hz, **Figure 2E**) and balanced synaptic currents (|*I_i_*/*I_e_*| > 0.75 for *v_a_* > 20 Hz, **Figure 2F**). Thus, the network displayed the features defining the spectrum of activity regimes over the entire range for 0.1 < *f* < 2. However, when increasing the excitatory and inhibitory recurrent synaptic weights by a larger factor (*f* > 2; see *f* = 5 in **Figure 2A**), the network generated states of dense activity throughout the entire range of afferent input rates (see yellow curves in **Figure 2C-F**), without displaying sparse activity states at low afferent input values, and showing a small range of variations of recurrent firing rate levels (from 15 Hz to 17 Hz, yellow curve in **Figure 2C**). These high recurrent synaptic weights led to strong recurrent amplification of afferent inputs, and the generated high recurrent activity levels led to negligible leak currents and to stabilization being achieved through dense recurrently-balanced dynamics (**Figure 2E,F**). For very low values of the *f* factor (we show the limit case of the absence of recurrent interactions defined by *f* = 0, dark purple curves in **Figure 2**), the population only displayed regimes dominated by afferent inputs 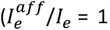, **Figure 2E**) and excitatory currents (|*I_i_*/*I_e_*| = 0, **Figure 2F**). This was due to negligible recurrent interactions which prevented the network from producing recurrently-balanced activity, therefore showing that a minimum amount of recurrent interactions was required to obtain the described spectrum of network states. Another factor influencing the strength of recurrent interactions was the amount of recurrent connections. Consistent with the need of moderate recurrent interactions, increasing the connectivity to make the network more densely recurrently connected that in the reference scenario (*p_conn_* > 20%) restricted the occurrence of AD-type activity to lower and lower afferent activity levels (see **Figure S3B**). This analysis emphasized the non-trivial nature of the appearance of the spectrum: it is a parameter-dependent emergent phenomenon at the network level (see Discussion).

**Figure 2.**
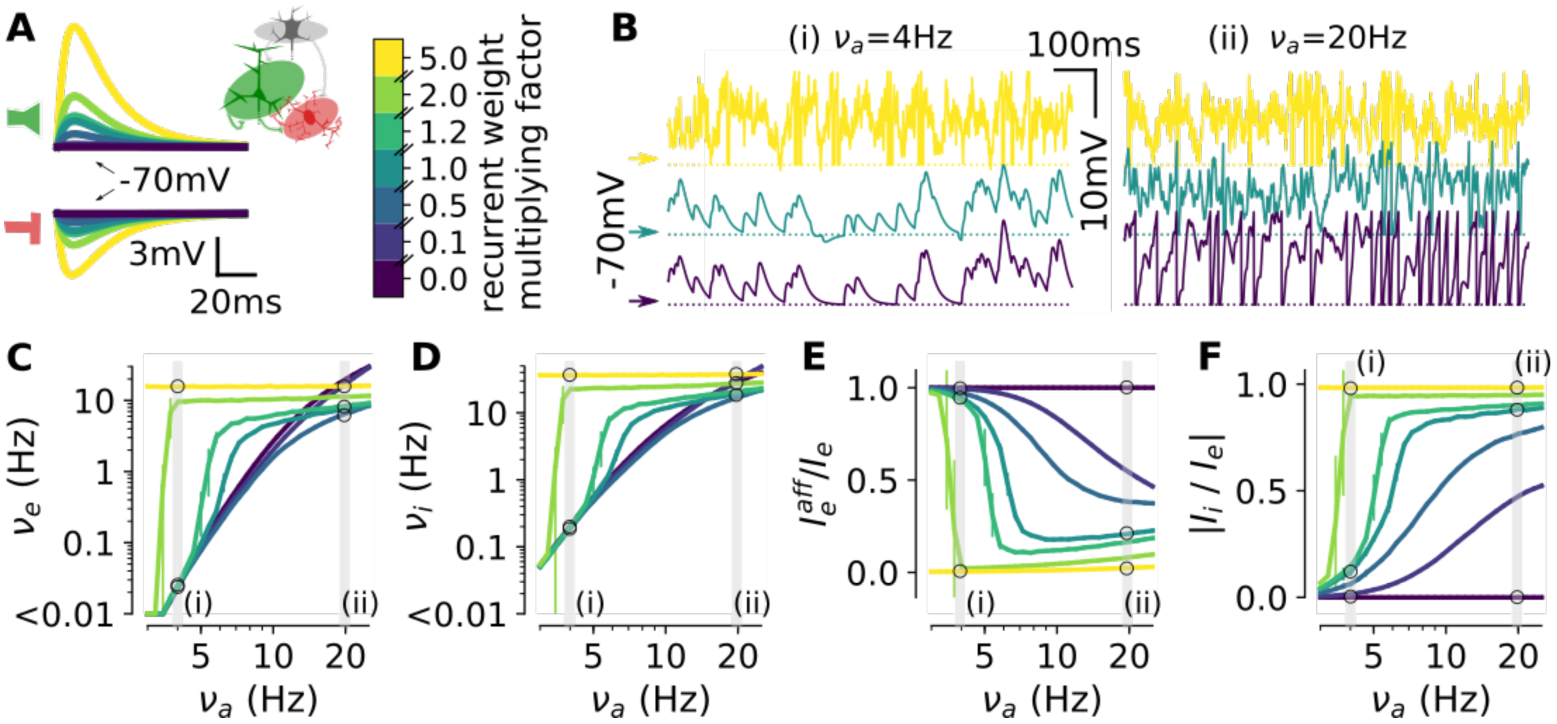
The emergence of the spectrum is conditioned to moderate strength of recurrent interactions. **(A)** We show the various post-synaptic deflections following an excitatory (top) and inhibitory (bottom) event as a function of the modulating factor for synaptic weights (color-coded, strong recurrent interactions in yellow (strength factor: 5), moderate in green (strength factor: 1) and absence of interaction in dark purple (strength factor: 0). **(B)** Sample traces of activity at low (left, (i), *v_a_* = 4 Hz) and high (right, (ii), *v_a_* = 20 Hz) levels of afferent activity for different strength of synaptic weights (same color code as in **A**). **(C-F)** Excitatory stationary firing rates (*v_e_* in **C**), inhibitory stationary firing rates (*v_i_* in **D**), fraction of afferent excitatory current 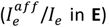, and inhibitory to excitatory current ratio (|*I_i_*/*_e_*| in **F**)as a function of the afferent activity (*v_a_*) for the seven different factors of recurrent synaptic weights. For a factor of two (green curve), the activity almost immediately jumps from a very low value (< 0.01 Hz) to a high activity level (*v_e_* ≳ 1 Hz) where synaptic activity is dominated by recurrent connectivity 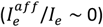 and it is balanced (|*Ii/*| *I_e_* 1). ~ For a factor of five (yellow curve), the activity is always high. Even very weak afferent inputs are translated into the balanced synaptic activity regime of high excitatory and inhibitory firing rates (see also panel **B(i)**). When negligible recurrent interactions are present (strength factor: 0), the activity remains purely dominated by its afferent excitation 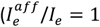 and *|I_i_/I_e_*|_=_0). We also show the lower bound configuration (*f* = 0.1). Above this factor, states dominated by recurrent activity at high afferent inputs 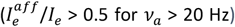 are observed. We represent the mean ± s.e.m (error bars) over n = 10 networks simulations. Non visible error bars correspond to variabilities smaller than the marker size. See **Figure S3** for the changes in other network parameters.

The other experimentally-driven constraints of our network implementation were much less critical for generating the spectrum of asynchronous dynamics. Varying the synaptic weights of the afferent input in the [-50 %, +50 %] range shifted the onset of the activity increase (in terms of *v_a_* level) but allowed for a set of asynchronous regimes across orders of magnitude (**Figure S3C**). Varying the inhibitory excitability by shifting the spiking threshold in the [−57, −52] mV range also did not affect the ability of the network to display a spectrum of activity regimes (**Figure S3D)**. This was also the case when varying the network size (**Figure S3E**) and the excitatory and inhibitory synaptic weights independently in the [-50 %, +50 %] range (**Figure S3F-G**). However, more extreme variations as the one listed below led to a recurrent architecture with a very strong excitatory-to-excitatory loop and produced saturated (*v_e_* = *v_i_* = 200 Hz) and highly synchronized (*SI* > 0.9) activity because of weak inhibition unable to prevent an excitatory runaway (Brunel, 2000). This happened for very low inhibitory excitabilities 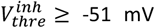 (**Figure S3E**), strong excitatory weights *Q_e_* ≥ 4 nS (**Figure S3F**), and weak inhibitory weights *Q_i_* ≤ 5 nS (**Figure S3G**). Low afferent input weights *Q_a_* ≤ 1 nS also prevented the appearance of the spectrum because only quiescent regimes (*v_e_* = *v_i_*) = 0 Hz) could be observed in the *v_a_* < 25 Hz range of afferent inputs (**Figure S3C**).

In sum, these results demonstrate that the ability of the network to generate a spectrum of states, in particular its ability to display the AD regime, depends crucially on having moderate strength of recurrent interactions.

### Introducing a disinhibitory circuit broadens the extent of the spectrum

To test the generality of the findings and its robustness to the inclusion of other physiologically-realistic circuit features, we considered a more complex network architecture encompassing a disinhibitory circuit, see **Figure 3A** and **Table 2**. The disinhibitory cells formed inhibitory synapses on inhibitory neurons. Because experimental evidence suggest weak inputs into disinhibitory cells from the local network (Jiang et al., 2015; Pfeffer et al., 2013), we assumed in the model that the disinhibitory cells received only excitatory afferent inputs. By lowering the excitability of the inhibitory population as a function of the afferent activity level, the disinhibitory activity allowed excitatory-dominated states to span higher ranges of firing rate values (up to *v_e_* = 58.3 ± 3.9 Hz for *v_a_* = 25 Hz, see **Figure 3B**) while remaining largely asynchronous (SI < 0.12, **Figure S3I**). The emergence of the spectrum of states was therefore robust to the inclusion of a disinhibitory circuit and its addition further strengthened and broadened the generation of the spectrum. As the model configuration inclusive of the disinhibitory circuit presumably provided a more realistic setting, we hereafter continued our analysis using the three-population model of **Figure 3A**.

**Figure 3.**
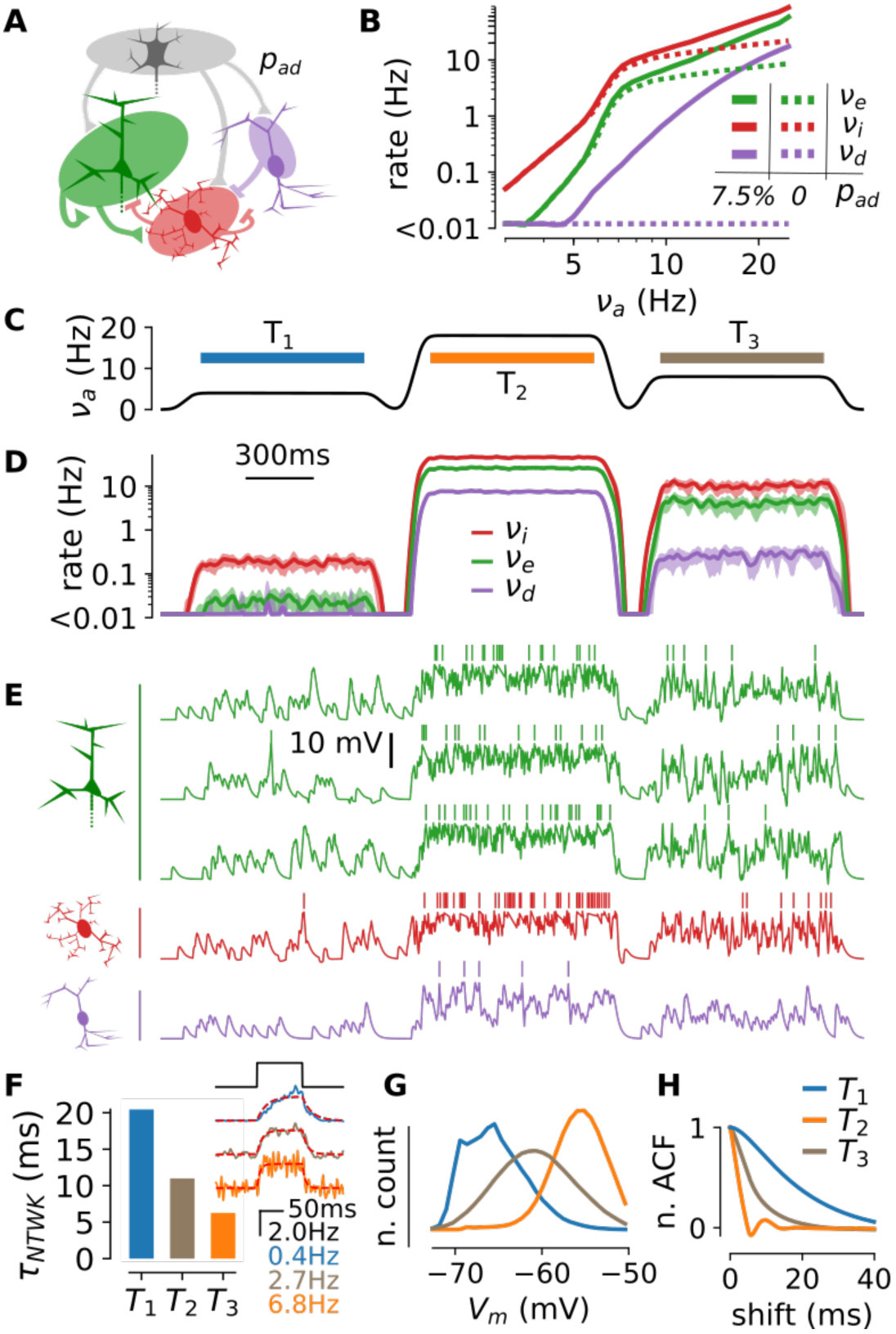
Dynamic modulation of network activity upon a time-varying afferent excitation in the three population model. **(A)** Schematic of the network model including the disinhibitory circuit. The parameter *p_ad_* corresponds to the connection probability between the afferent and disinhibitory population. Model parameters can be found in **Table 2**. **(B)** Stationary co-modulations of the excitatory (*v_e_*, green), inhibitory (*v_i_*, red), and disinhibitory (*v_d_*, purple) rates in absence (*p_ad_* = 0, dashed line, reproduced from **Figure 1B**) and in presence of a disinhibitory circuit (solid line, *p_ad_* = 7.5%). **(C-E)** Network dynamics in response to a time-varying input. **(C)** Waveform for the afferent excitation. **(D)** Temporal evolution of the instantaneous firing rates (binned in 2 ms window and smoothed with a 10 ms-wide Gaussian filter) of the excitatory (*v_e_*, green), inhibitory (*v_i_*, red), and disinhibitory (*v_d_*, purple) populations. Mean ± s.e.m over n = 10 trials. **(E)** Sample membrane potential traces in a trial (three excitatory cells in green, an inhibitory cell in red, and a disinhibitory cell in purple). Note that, to highlight mean depolarization levels, the artificial reset and refractory mechanism has been hidden by blanking the 10 ms following each spike emission. (**F**) Network time constants *τ_NTWK_* for the three different levels of afferent activity considered in **C** (*v_a_* = 4 Hz in blue, *v_a_* = 18 Hz in orange, *v_a_* = 8 Hz in brown). The time constant was determined by stimulating the network with a 100 ms-long step input of afferent activity of 2 Hz (black curve in the inset) and fitting the trial-average responses with an exponential rise-and-decay function (red dashed curves, see Methods for details). We show the average over 100 stimulus repetitions of the network responses in the inset. **(G-H)** Electrophysiological signature in the three intervals highlighted in **C:** *T*_1_ (blue), *T*_2_ (orange), and *T*_3_ (brown). Data were obtained pooling together the *V_m_* after blanking spikes over 100 excitatory neurons in each interval for a single network simulation. **(G)** Pooled membrane potential histograms for the three different stimulation periods. (**H**) Pooled normalized autocorrelation functions.

**Table 2.**
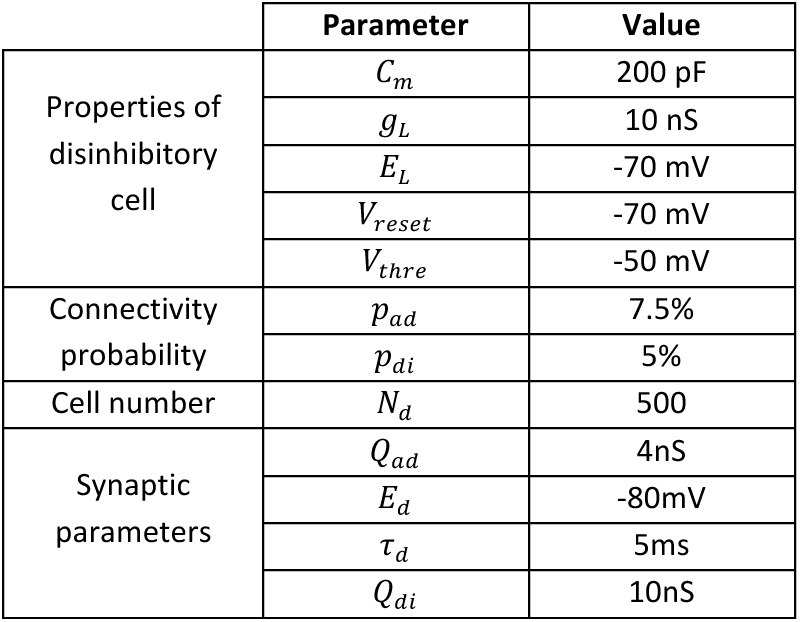
Additional parameters to construct the three
population model (excitation/inhibition/disinhibition).

### Dynamic modulation of the network regime upon time-varying afferent excitation

After analyzing the stationary behavior of the network model, we studied whether it could generate a spectrum of states in the presence of time-varying inputs. In particular, given that in awake cortical data different states can persist for time scales of < 1 s (McGinley et al., 2015a), we focused on studying network dynamics when inputs were stationary for some hundreds of milliseconds. In **Figure 3C**, we stimulated the three population model with a time-varying waveform made of three 900 ms-long plateaus of presynaptic activity at low (*v_a_* = 4 Hz, *T*_1_ period, blue interval), high (*v_a_* = 18 Hz, *T*_2_ period, orange interval), and intermediate (*v_a_* = 8 Hz, *T*_3_ period, grey interval) levels. **Figure 3D** shows the temporal evolution of the firing rates (averaged over n = 10 trials) and **Figure 3E** shows the *V_m_* dynamics in the three cellular populations included in the model in a single trial. We observed dynamic modulation of the firing rate with time scales to reach stationary behavior which were similarly fast across the different cell types (red, green, and purple in **Figure 3D**). Indeed, when computing the relaxation time of the network (τ_*NTWK*_, estimated by fitting the response to a short step of afferent activity, see **Figure 3F**), we found network time constants between 4 ms and 20 ms (with a monotonic dependence on the level of ongoing activity, as predicted theoretically (Destexhe et al., 2003; van Vreeswijk and Sompolinsky, 1996)). For time scales longer than few hundred ms, the network dynamics can thus be considered as stationary. Consequently, the characterization described above for stationary states (**Figure 1** and **Figure 3B**) also held when the analysis was restricted to the three separate windows *T*_1_, *T*_2_ and *T*_3_ (see **Figure 3C-E**). For example, in the first period (*T*_1_), the spiking activity was temporally sparse (*v_e_* = 0.02 ± 0.01 Hz), the average *V_m_* value was hyperpolarized (*μ_V_* = −65.3 ± 0.1 mV), the *V_m_* fluctuations were skewed (*γ_V_*= 0.55 ± 0.01, blue curve in **Figure 3G**), and the autocorrelation function displayed large *τ_V_* values (blue curve **Figure 3H**, leading to an estimate of *τ_V_* = 17.8 ± 0.3 ms). In the second period (*T*_2_), the afferent input increased spiking activity by four orders of magnitude (*v_e_*= 25.9 ± 0.6 Hz) and it shifted the *V_m_* dynamics to depolarized Gaussian fluctuations (orange curve in **Figure 3G**, *μ_V_* = −55.9 ± 0.1 mV, *R*^2^ of a Gaussian fitting after blanking spikes: 0.99 ± 0.01) characterized by a narrowly extended autocorrelation function (orange curve in **Figure 3H**, leading to an estimate of *τ_V_* = 2.3 ± 0.1 ms). The last segment (*T*_3_) was an intermediate situation that produced excitatory activity at *v_e_* = 4.2 ± 0.3 Hz associated to moderate depolarization with high variance fluctuations (grey curve in **Figure 3G**, *μ_V_* = −60.8 ± 0.2 mV and *σ_V_* = 4.3 ± 0.1 mV) and an intermediate autocorrelation time (*τ_V_* = 6.8 ± 0.3 ms from the orange curve in **Figure 3H**). The possibility to identify a given network state from the spiking activity and from the *V_m_* fluctuations at the sub-second time scale prompted us to test the model’s prediction with experimental electrophysiological recordings.

### In the somatosensory cortex of awake mice diverse epochs of non-rhythmic activity have electrophysiological signatures predicted by the model

How does neural activity in real cortical circuits compare with the activity regimes observed in the model? To address this question, we performed intracellular patch-clamp recordings from layer 2/3 neurons of the somatosensory cortex of awake mice (n = 22 cells in N = 8 animals) during spontaneous activities (**Figure 4**). These recordings (**Figure 4A**) showed fluctuations in the membrane potential of recorded cell between rhythmic and asynchronous dynamics as described in previous reports (Crochet and Petersen, 2006; Poulet and Petersen, 2008), see also **Figure 4B** for examples of individual time epochs. Because our focus was asynchronous cortical dynamics, we introduced a threshold in the low frequency power of the *V_m_* recordings, which we called “rhythmicity threshold” (see **Methods**). We classified as rhythmic periods all the time stretches for which the *V_m_* power exceeded this threshold (Figure 4A) and we considered for further analyses only the epochs of network activity with *V_m_* power below this threshold (classified as “non-rhythmic”, see **Figure 4A**; note that the results were robust to variations of this arbitrary threshold, see **Figure S4D**).

**Figure 4.**
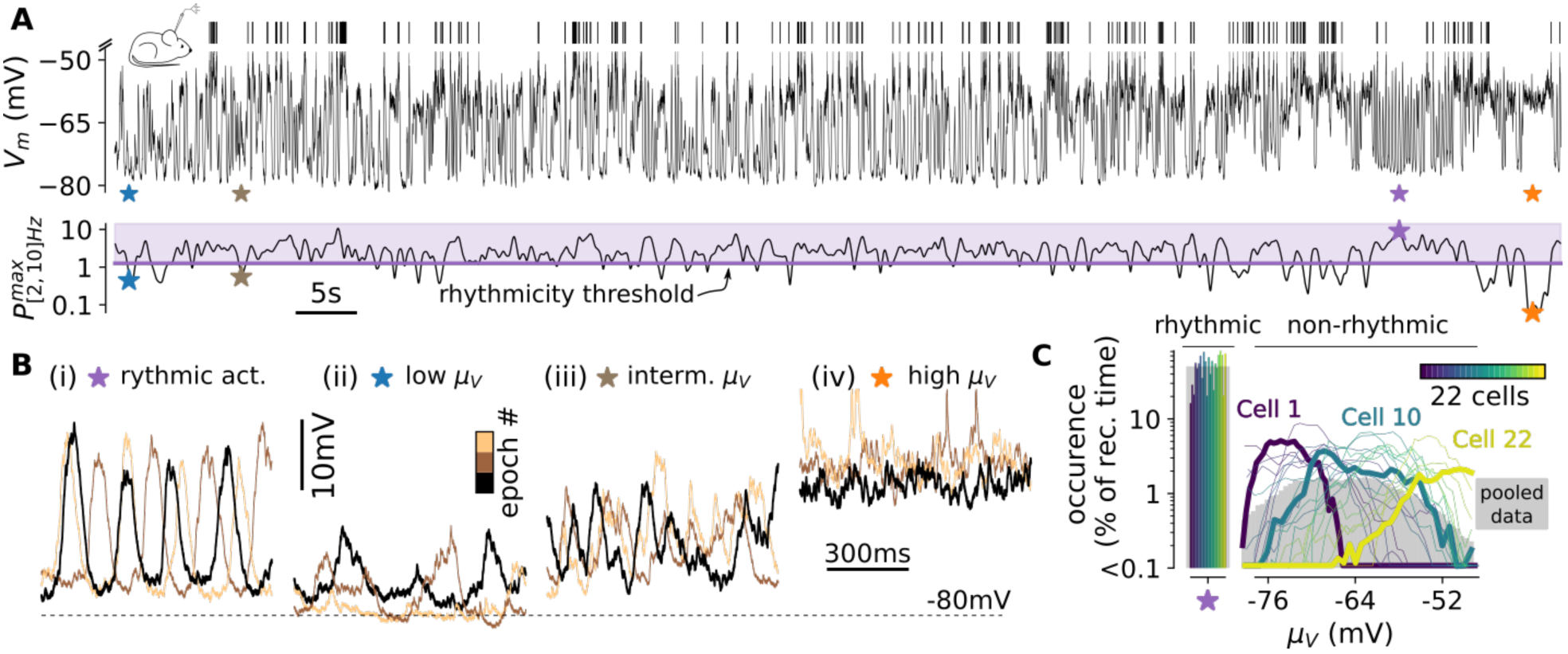
In the barrel cortex of awake mice, non-rhythmic activity is associated with various membrane depolarization levels. **(A)** Intracellular recordings of *V_m_* fluctuations (top) during spontaneous activity and maximum power of *V_m_* in the [2, 10] Hz band (bottom). Within this 103 s sample, we highlight the occurrence of three periods classified as non-rhythmic epochs (blue, brown, and orange stars) and one rhythmic epoch (purple star). Note the 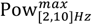 index being below (for the three non-rhythmic events) and above (for the rhythmic event) the rhythmicity threshold. **(B)** Three *V_m_* samples classified as (i) rhythmic epochs, (ii) low *μ_V_* epochs: *μ_V_* < −70mV, (iii) intermediate *μ_V_* epochs: *μ_V_* ∈ [−70, −60] mV and (iv) high *μ_V_* epochs: *μ_V_* > −60 mV. The black traces correspond to the samples highlighted in **A**, the two other samples (copper colors) were extracted from the same intracellular recording. **(C**) Fraction of occurrence of the rhythmic activity epochs together with the non-rhythmic epochs at their respective levels of mean depolarization *μ_V_*. Single cell recordings have been sorted with respect to their average level of non-rhythmic activity *μ_V_* and color-coded accordingly. We highlight three cells (see main text): Cell 1, Cell 22, and Cell 10 (shown in **A** and **B**). The plain grey area represents the dataset after pooling together all *V_m_* recordings (n = 22 cells). Note that the fraction of occurrence of rhythmic activity in the pooled data corresponds to 50% as a consequence of the definition of the rhythmicity threshold (see **Methods**).

We then divided the stretches of non-rhythmic activity into a series of 500 ms-long epochs (partly overlapping, and taken with sliding windows whose center was moved in steps of 25 ms). Each epoch was considered a possible different state. We chose this epoch length as it offered a good compromise between the following two constraints: 1) it was short enough to enable the identification of specific states of wakefulness (this amounts to choosing T < 1 s, because a time scale longer than 1 s would likely merge different network states, see (McGinley et al., 2015a)) and 2) it was long enough to average synaptically-driven *V_m_* dynamics and to analyze network activity beyond its own relaxation time constant (i.e. T ≫ 10 ms, see the previous section for the model’s properties and (Reinhold et al., 2015) for a concordant experimental estimate in the mouse visual cortex). We then examined in detail the properties of neural activity in each such epoch. Similarly to previous findings in the auditory (McGinley et al., 2015b) and visual (Reimer et al., 2014) cortices of awake behaving mice, we found non-rhythmic epochs of network activity in the primary somatosensory cortex that showed various levels of mean membrane depolarization *μ_V_*. **Figure 4B** shows representative membrane potential epochs and their fraction of occurrence at the various *μ_V_* levels over single cells (color-coded in **Figure 4C**) and over the ensemble data (grey area in **Figure 4C**). Few cells (n = 3 out of 22, for example “Cell 10” shown in **Figure 4A,B**) displayed non-rhythmic activity over a wide range of *μ_V_* levels (> 20 mV). The majority of cells displayed non-rhythmic activity in a narrower range of depolarization levels *μ_V_* (for the remaining n = 19 out of 22 the extent of *μ_V_* was 10.8 ± 4.1 mV; e.g. “Cell 1” corresponded to a recording exhibiting only hyperpolarized non-rhythmic activity and “Cell 22” corresponded to a recording exhibiting mostly depolarized non-rhythmic activity, see **Figure 4C**). This limited variability might have partly been due to the relatively short durations of our *V_m_* samples that resulted from the strict criteria we used to define stable *V_m_* recordings (enabling us to perform the analysis on an absolute *μ_V_* scale, see **Methods**).

One central prediction of the model was the occurrence of a range of different states at various mean depolarization levels *μ_V_* with mean spiking activity spanning over 3 - 4 orders of magnitude (see inset in **Figure 5A**). This prediction was confirmed in our experimental data: hyperpolarized epochs displayed firing rates below 0.1 Hz, while depolarized epochs exhibited firing activity in the 10 Hz range (see **Figure 5A**, in the right inset we show representative epochs exhibiting spikes at low, intermediate, and high *μ_V_* levels). The wide range of firing rates (suggestive of exponential amplification of spiking activity) across the non-rhythmic states of wakefulness with different mean depolarization levels *μ_V_* was further confirmed by extracellular recordings (see **Figure S5**). Indeed, we combined the previously described intracellular recordings with extracellular recordings of the multiunit activity in layer 2/3 (N = 4 mice, n = 14 cells, see **Methods**). We found that the logarithm of the mean multi-unit activity within non-rhythmic epochs exhibited a robust linear correlation with the depolarization levels *μ_V_* (correlation coefficient c = 0.5, one-tailed permutation test: p < 1e-5, see **Figure S5**). This suggests that the wide range of rates predicted by the model was observed not only at the single neuron level but also at the mass circuit activity level, as expected by the theoretical model.

**Figure 5.**
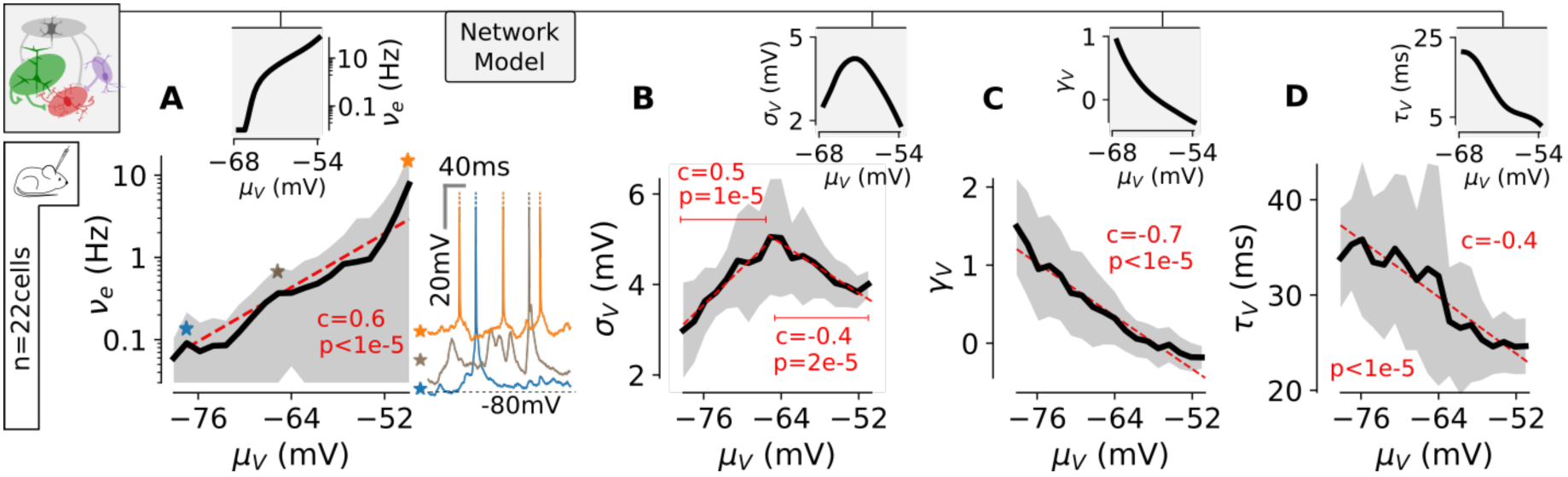
Various levels along the theoretical spectrum predict the electrophysiological signature of the different non-rhythmic epochs of wakefulness in mouse barrel cortex. **(A)** Spiking probability (*v_e_* in Hz) of intracellularly recorded layer 2/3 pyramidal cells within *μ_V_*-classified epochs. The red dashed line is a linear regression between *μ_V_* and *log*_10_(*v_e_*) (see **Methods**). In the right inset, we show 300 ms-long sample epochs displaying spikes for three levels of *μ_V_* (blue, brown, and orange stars in main plot). See also **Figure S5** for extracellular recordings of population spiking activity. In the main plots, we show the co-modulation between the depolarization level and *μ_V_* (at a given *μ_V_* level, mean ± s.e.m. over the cells displaying that specific *μ_V_* level, see **Figure 4C**). The correlation coefficients and the p-value of a one-tailed permutation test (see **Methods**) are reported. In top inset, we show the predictions of the network model. **(B)** Co-modulation between the depolarization level *μ_V_* and the standard deviation of the fluctuations *σ_V_*. Note that the linear regression has been split into two segments (depicted in red) to test the significance of the non-monotonic relationship. **(C)** Co-modulation between *μ_V_* and the skewness *γ_V_* of the *V_m_* distribution. **(D)** Co-modulation between *μ_V_* and the autocorrelation time *τ_V_* of the *V_m_* fluctuations.

A more detailed characterization of the correspondence between model predictions and *in vivo* dynamics was obtained by analyzing how the features of *V_m_* in each epoch depended on the mean membrane depolarization *μ_V_* level in the same epoch. We measured in real data the following feature of *V_m_* fluctuations that were previously described in our model: 1) the standard deviation, *σ_V_*; 2) the skewness of the *V_m_* distribution, *γ_V_*; 3) the speed of the *V_m_* fluctuations quantified by the autocorrelation time, *τ_V_* (see **Methods**). The network model predicted that: 1) the *σ_V_* -*μ_V_* relationship should be non-monotonic with a peak in the intermediate *μ_V_* range (inset of **Figure 5B**); 2) the *γ_V_* -*μ_V_* relationship should start from strongly positively skewed values (*γ_V_* ~ 1) and monotonically decrease with *μ_V_* (inset of **Figure 5C**); 3) the *τ_V_* -*μ_V_* relationship should be monotonically decreasing with a near 15 ms drop in *τ_V_* (inset of **Figure 5D**). Remarkably, we found all those features in the *V_m_* fluctuations properties of our experimental recordings (**Figure 5B-D**). Moreover, those relationships were found highly significant as the null hypothesis of a zero-slope relationship corresponded to very low probabilities (p < 5e-5 for all relationships, see the individual p-values in **Figure 5B-D**). The model prediction of a transition toward Gaussian fluctuations at high *μ_V_* (**Figure 3F**) was also found to hold on real data: we fitted the pooled distributions with a Gaussian curve (see **Methods**) and the coefficient of determination was found to be *R*^2^ = 0.99 ± 0.01 above *μ_V_* = −60 mV compared to *R*^2^ = 0.96 ± 0.04 below *μ_V_* = −60 mV (n = 55 *μ_V_*-defined distributions across 13 cells for *μ_V_* > −60 mV and n = 129 *μ_V_*-defined distributions across the 22 cells for *μ_V_* ≤ −60 mV, p = 3.2e-5, unpaired t-test). Taken together, these results show that the spectrum of activity in the theoretical model predicted, in real data, the electrophysiological features of the non-rhythmic epochs as a function of their *μ_V_* level both in terms of spiking activity and *V_m_* fluctuations.

### Activity levels along the spectrum are characterized by different computational properties

The *in vivo* recordings displayed in **Figure 4** showed that networks transiently settle at various points along the spectrum of asynchronous dynamics. Does the shift between activity states within the spectrum affect the capabilities of the circuit to encode afferent information? To investigate how the transition between asynchronous activity regimes may shape the computational properties of the network, we designed two types of afferent stimulus sets that we fed to the model, both in the AD regime and in the RD regime (**Figure 6**).

**Figure 6.**
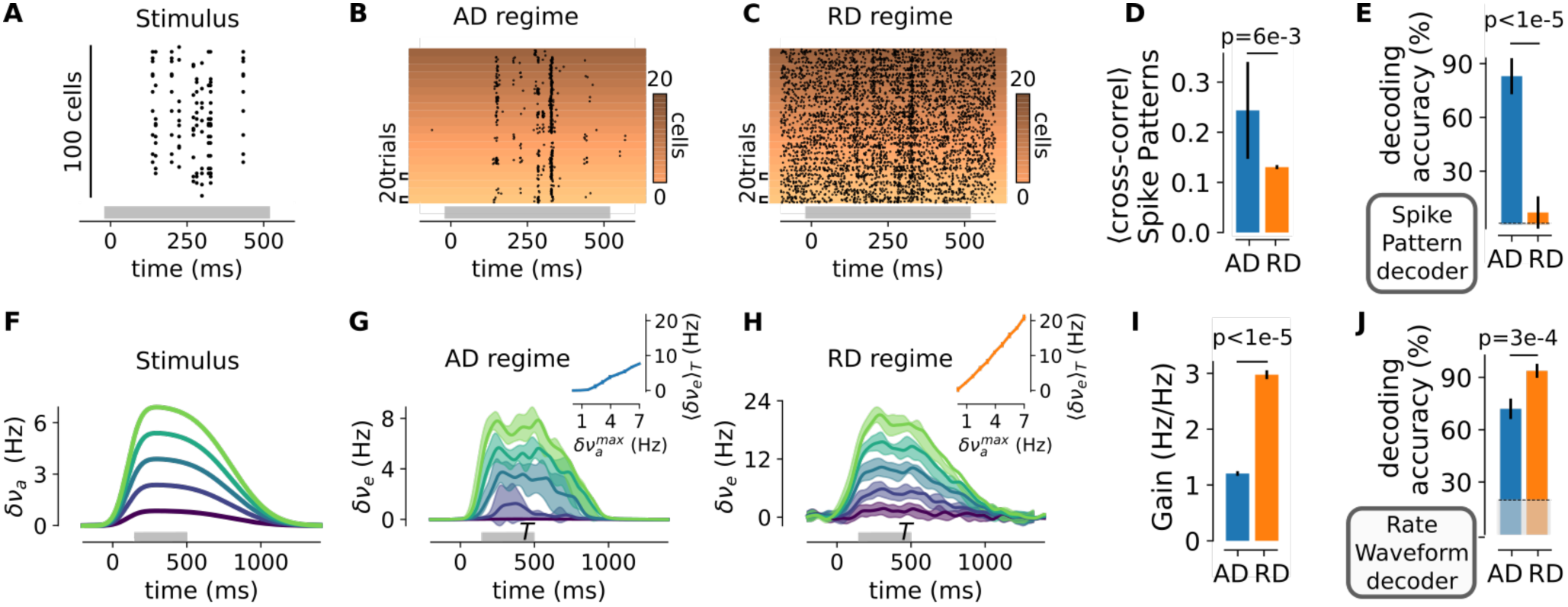
Distinct computational properties along the activity spectrum: the AD regime enables the precise encoding of complex patterns of presynaptic activity while the RD regime exhibits high population responsiveness to afferent inputs. **(A)** Representative example of a presynaptic activity pattern which corresponds to ten activations of different groups of ten synchronously spiking units (randomly picked within the one hundred cells of the presynaptic population) in a 500 ms window (see **Methods**). **(B)** Spiking response of a sub-network of neurons (20 cells) across 20 trials in the AD regime. The y-axis indexes both the neuron identity (color-coded) and the trial number (vertical extent on a given color level). **(C)** Same than **B** for the RD regime. **(D)** Mean cross-correlation of the output spiking patterns across realizations for a given input pattern (mean ± s.e.m over 10 input patterns, for each input pattern we computed the mean cross-correlation across all pairs of observations of the 20 realizations, two-sided Student’s *t*-test). **(E)** Performance in decoding the pattern identity from the sub-network spiking patterns with a nearest-neighbor classifier (see Methods). The mean accuracy ± s.e.m. over ten patterns of ten test trials each is shown (two-sided Student’s *t*-test). The thin dashed line indicates chance level (from 10 patterns and 10 onsets: 1%). **(F)** The model network is fed with a stimulus whose firing rate envelope is of varying maximum amplitudes stimulus 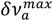 (amplitude values are color-coded). **(G)** Mean and standard deviations over n = 10 trials of the increase in excitatory population activity 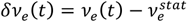 in the AD regime. The stationary activity level 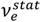 was discarded and the rates were smoothed with a 30 ms Gaussian filter. In the inset, the response average in the time window *T* (highlighted by a grey bar on the time axis) as a function of the maximum amplitude of the stimulus 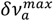 is shown. **(H)** Same than **F** for the RD regime. **(I)** Slope of the relationship between 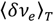 and 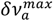 (mean +/- s.e.m over n=10 trials, statistical analysis: two-sided Student’s *t*-test). **(J)** Decoding the sub-network rate waveform with a nearest-neighbor classifier (see **Methods**). The thin dashed line indicates chance level (from the 5 waveforms shown in **F**: 20 %). The mean accuracy ± s.e.m. over five patterns of ten test trials each is shown (two-sided Student’s *t*-test).

The first stimulus set mimicked the precise spatiotemporal patterns often evoked by sensory stimuli (Foffani et al. 2009; Luczak et al., 2015; Panzeri et al., 2010; Petersen et al., 2008; Urbain et al., 2015). It corresponded to a pattern of sequential presynaptic co-activations, distributed over a 500 ms time window and it targeted a subset of 100 neurons within the full network (maintaining the connectivity probability and synaptic weights of the background activity, see **Methods**). We show in **Figure 6A** an example of such an afferent pattern. **Figure 6B** and **6C** show the response in the targeted sub-network over different trials (where the realizations of background activity vary) for the AD and RD regimes, respectively. The network activity across trials was highly structured by the stimulus in the AD regime (**Figure 6B**), while the stimulus-evoked response was less reliable and more diluted in the RD regime (**Figure 6C**). We generated various random realizations of such afferent patterns (see other examples in **Figure S6A**) and analyzed the reliability of the various responses across trials using a scalar metric for multi-unit activity (van Rossum 2001, with a temporal sensitivity of 5 ms, see **Methods**). We found that the trial-to-trial cross-correlation between the output spiking responses and the presented afferent pattern was significantly larger in the AD than in the RD regime (p = 6e-3, paired t-test, see **Figure 6D**). By including the distance of the above metric in a nearest-neighbor classifier, we constructed a decoder retrieving both the pattern identity and the stimulus onset from the output spiking activity of the target population (see **Methods**). We used this classifier to analyze whether the ability of the AD regime to generate reliable output patterns (reported in **Figure 6D**) would lead to a robust joint decoding of both the identity and onset timing of the afferent input pattern. We found that these spatiotemporal features of the afferent input pattern were faithfully encoded by the activity of the target network in the AD regime (accuracy for the joint decoding of both afferent pattern identity and onset: 83.0 ± 10.1 %, **Figure 6E**). In contrast, in the RD regime the decoding accuracy remained close to chance level (7.0 ± 9.0 %; **Figure 6E)**. The explanation for this difference can be found in the drastically different levels of activity in the two regimes (*v_e_* = 0.02± 0.01 Hz for the AD regime compared to *v_e_* = 25.9 ± 0.6 Hz for RD). In the AD regime, the patterned structure of the input strongly constrained the spiking activity of the population (as the stimulus-evoked spikes represented 92.4 ± 4.4 % of the overall activity), therefore leading to a reliable encoding of the input identity. In the RD regime, the stimulus-evoked spiking was confounded by the strong ongoing dynamics in single trials in the RD regime (stimulus-evoked activity only represented 6.7 ± 5.7 % of the overall activity) therefore impeding a reliable decoding of activity patterns.

We then fed the network with waveforms of afferent activity at various amplitudes targeting the entire network without any spatiotemporal structure within the stationary period of the afferent waveform (see Methods and **Figure 6F**). We decoded the level of afferent activity from the sum of excitatory population activity within the recurrent network (**Figure 6G-H**, for the AD and RD regimes, respectively). Classically (Murphy and Miller, 2009; Tsodyks and Sejnowski, 1995; Van Vreeswijk and Sompolinsky, 1996), in the RD regime, the network showed a linear response of high gain (**Figure 6I** and inset of **Figure 6H**) and the response waveforms accurately represented the level of the afferent input waveforms (see **Figure 6H**). We found that, however, this was not the case in the AD regime. In this regime, the population response exhibited a weak amplification of the input signal (note the much lower gain in **Figure 6I**) and it failed to accurately follow the input (single trial responses in the AD regime had significantly lower cross correlations with the input waveform: 0.81 ± 0.26 for AD *vs* 0.87 ± 0.23 for RD, p = 4.4e-3, two-tailed Student’s *t*-test). When decoding the input signal from the single-trial time-varying rate of a small populations of the network (100 excitatory neurons, a population size equal to that used above for the decoding of the precise afferent spatiotemporal patterns), we now observed a higher decoding accuracy in the RD regime than in the AD regime (**Figure 6J**, nearest-neighbor decoder combined with a rate waveform metric, see **Methods**), suggesting that the RD regime favors the reliable encoding of the overall strength of the afferent activity thanks to its high amplification properties.

## Discussion

Our study reports a novel emergent feature of recurrent dynamics in spiking network models: a spectrum of asynchronous activity states in which firing activity spans orders of magnitude and in which the predominance of the synaptic activity shifts from the afferent input component (AD) to the recurrent input component (RD). Importantly, the continuous set of network states predicted by the model matches qualitatively and quantitatively the set of non-rhythmic cortical states observed in awake rodents. Our model predicts that each regime has marked electrophysiological signatures, and the model predictions were validated by experimental recordings of asynchronous activities in superficial layers of the somatosensory cortex in awake mice. Moreover, we found that, under specific biophysical constraints (discussed below), two qualitatively different computational properties could coexist within the same network: the reliable encoding of complex presynaptic activity patterns in the AD regime together with the fast and high-gain response properties associated to balanced recurrent amplification in the RD regime.

Using rate-based models of cortical dynamics, previous theoretical analysis suggested that recurrent networks can be made to operate in afferent-driven regime and recurrent-driven regime (Ahmadian et al., 2013; Rubin et al., 2015). However, this seminal work left important questions unanswered. Being a phenomenological rate model, it could neither investigate the detailed biophysical mechanisms behind the creation and coexistence of these regimes in the same network nor reveal the computational advantages in terms of information coding of each state resulting from their spiking dynamics. Finally, this previous work was proposed as a model for multi-input integration in sensory cortices and did not investigate whether such a theoretical picture could be used to explain, in quantitative terms, the electrophysiological signature of neocortical dynamics across the different states of wakefulness. The present study developed those aspects through the combination of biophysically plausible spiking network modeling and experimental recordings in awake rodents.

### Key features of the model shed light on the biophysical circuit-level mechanisms for the emergence of the spectrum

The present study detailed the biophysical mechanisms as well as the necessary conditions for the emergence of spectrum of states. Our work showed that the observed spectrum of asynchronous regimes and their functional diversity is an emergent dynamical property of spiking networks, because it relies on the interactions between the different mechanisms characterizing such systems: synaptically-mediated single cell integration, spike-and-reset mechanisms, and leak repolarization currents (see **Figure 1**). Unlike previous analysis where afferent synaptic currents were described by stochastic processes only constrained by a mean and a variance (Brunel, 2000; Renart et al., 2004; van Vreeswijk and Sompolinsky, 1996), we explicitly modeled afferent activity as a shot noise process producing post-synaptic events of excitatory currents. At the single cell level, this feature was crucial to produce skewed membrane potential distribution (DeWeese and Zador, 2006; Richardson et al., 2010; Tan et al., 2014). At the network level, it enabled the emergence of the AD regime. Crucial to our model was also the presence of conductance-based interactions. This feature of the model allowed synaptic efficacy to be high at low levels of activity while being strongly dampened at higher level (Kuhn et al., 2004). Other types of activity-dependent modulation of synaptic transmission such as short-time depression (Tsodyks and Markram, 1997) or synaptic disfacilitation (Urban-Ciecko et al., 2015) may also contribute to this mechanism. On a dynamical perspective, this property constrained uncontrolled increase of the *V_m_* fluctuations upon a 2-3 orders of magnitude raise in recurrent activity and helped maintaining stable asynchronous dynamics over the large range of firing rates. Note however, that, although this feature of single cell integration is a necessary mechanism for the emergence of the spectrum, it is not a sufficient condition and the non-monotonic *σ_V_*-*μ_V_* relationship is not generally observed in the network model (in the “strong recurrence” network of **Figure 2**, although individual cell contain this single cell mechanism, the emergent dynamics at the network level does not display the non-monotonic relationship because it only displays RD regimes).

The key variable governing network state modulation in the model was the level of afferent excitation (**Figure 1** and **3**). In accordance with such a dependency, shifts in network state in the cortex have been observed to be controlled by thalamic excitation (Poulet et al., 2012). Network state modulation has also been shown to be regulated by the activity of other subcortical structures (Reimer et al. 2016; Zagha and McCormick, 2015). Whether such contribution is mediated by a net increase in afferent excitatory input (i.e. by the effect described in **Figure 1, 3**), by the neuromodulation of effective synaptic weights (i.e. by the effect described in **Figure 2**), or by any other mechanism remains to be established.

Another critical network setting to obtain the described spectrum of regimes was a moderate strength of recurrent interactions (**Figure 2**). It is thus important to understand whether this assumption is valid in light of the experimental evidence accumulated so far. Reducing the strength of recurrent interaction to single numbers is, however, difficult given the high heterogeneity of excitatory and inhibitory cells found in the neocortex (heterogeneity which is not entirely replicated in our simplified model (Jiang et al., 2015; Markram et al., 2015)), and given the area, layer, and species specificities that are often experimentally observed. This complexity notwithstanding, we restrict our discussion here to mouse experimental data on the superficial cortical layers (layer 2/3, where our experiments were performed). Unitary post-synaptic potentials observed in slice recordings (maximum amplitudes below 2 mV (Jiang et al., 2015; Lefort et al., 2009; Markram et al., 2015)) are in agreement with the conditions of “moderate weights” that we used in our model for both excitatory and inhibitory synaptic transmission (at −70 mV, our model gives maximal amplitudes of *δV*=2.1 mV for excitatory synapses and *δV* =-1.4 mV for inhibitory synapses, see **Figure 1A**). Moreover, from local measurements of excitatory projections in adult rodent cortex (Jiang et al., 2015), recurrent excitatory connections seem to match the “sparse connectivity” requirement with connectivity probabilities below 10 % (note that measurements in the juvenile cortex point toward slightly higher values in the 10-20% range (Lefort et al., 2009; Markram et al., 2015), see also (Barth et al., 2016)). In contrast, local measurements of inhibitory projections in adult mice show high (> 30 %) connectivity probabilities, in particular when targeting interneuronal sub-populations (Jiang et al., 2015). However, two phenomena might decrease the effect of this experimental finding when considering our simplified excitatory/inhibitory network framework. First, given the spatial attenuation of those connectivity probabilities (Jiang et al., 2015), a strong reduction is expected when considering the spatial scale studied here (5000 neurons), corresponding to ~ 0.2 mm^2^ of cortical tissue (Markram et al., 2015). Second, connectivity values and connectivity patterns of inhibitory projections largely vary depending on the type of source and target neurons (Pfeffer et al., 2013). For example, while PV cells show a high connectivity with excitatory neurons, other types of interneurons (e.g., the vasoactive intestinal peptide–expressing (VIP) cells) show low level of connectivity with excitatory neurons. This high heterogeneity across interneuronal subtypes may thus result in moderate average connectivity values for inhibitory projections, despite some interneuronal classes showing high connectivity with specific targets. Altogether, although previous experimental observations provide evidence in support of our model setting, the extent to which moderate strength of recurrent interactions in the neocortex can be extended across different cortices and animal species remains to be determined. This is even more true considering that the strength of recurrent connectivity may vary over time (e.g., at different developmental stages and in an activity-based manner).

Regarding cell heterogeneity and cell type-specific effects, it is interesting to note that the activity of one main class of disinhibitory interneurons, the VIP interneurons, was observed to be strongly modulated across behavioral states (Lee et al., 2013; Pi et al., 2013). The VIP population has been shown to control the level of local recurrent activity (Jackson et al., 2016; Pi et al., 2013), the motor integration in the primary somatosensory cortex (Lee et al., 2013) and the gain of cortical responses (Fu et al., 2014). Similarly, in the model, the inclusion of a disinhibitory circuit broadened the extent of the spectrum of regimes by enabling high activity states in the network (**Figure 3B**) which were associated to strong recurrent amplification properties (**Figure 6G-H**).

### Electrophysiological signature of non-rhythmic network regimes: model *vs* experiments

One important feature of our theoretical description is that it makes clear predictions about how several major experimentally measurable quantities change across states. Generating testable prediction is key for using the model to understand which circuit mechanisms are at work in the real cerebral cortex, and for understanding how to further refine the realism and predictive power of the model. In our case, the model generated four predictions (derived both from the numerical analysis of the model, see main text, and from its analytical study, see **Supplementary Information**) that were non-trivially expected to happen in any model. This non-trivial aspect is visible on **Figure 2**, the “strong recurrence” network (yellow curves) contains the same biophysical mechanisms than the “moderate recurrence” network, but does not lead to the same electrophysiological signature because of the parameter dependency of the emergent dynamics. We could therefore analyze such signature on cortical data and thus give insights about cortical mechanisms and circuit function.

A major prediction of our model, confirmed by the cortical data, was the presence of a strongly non-monotonic relationship between *σ_V_* and *μ_V_* (**Figure 5B**) as neural activity increases across states in the spectrum (**Figure 5A**). This relationship goes against a simple expectation (Campbell’s theorem, see (Daley and Vere-Jones, 2003)) which is derived from considering *V_m_* dynamics as the result of summed excitatory and inhibitory processes (Kuhn et al., 2004). Importantly, experimental measurements that we performed in the somatosensory cortex of awake mice (**Figure 5B**), as well as earlier data (McGinley et al., 2015b; Polack et al., 2013), confirmed the predictions of the model and exhibited a non-monotonic *σ_V_* - *μ_V_* relationship with a clear decay at high *μ_V_*, suggesting the presence of this key model mechanism in real cortical circuits. Our model also predicted that the autocorrelation time of the *V_m_* fluctuations *τ_V_* decreases with the mean membrane potential value *μ_V_*. This feature, which we also found to be present in cortical data (we observed a linear decrease of *τ_V_* with *μ_V_*; **Figure 5D**), in terms of mechanisms is likely due to the fact that single neuron integration is faster upon an increase of recurrent activity (Destexhe and Paré, 1999). From the functional point of view, this feature is crucial to enable the non-trivial dynamical property of displaying fast responses in the RD regime (**Figure 6H**), resulting in the ability of potentially tracking variations in the input at time scales of few milliseconds. The model predicted that *γ_V_* is reduced as the frequency of post-synaptic events increases and therefore makes the distribution closer to its “diffusion approximation”, i.e. a Gaussian distribution (analogous to a “large number limit” where statistical moments beyond second order tend to vanish, see (Daley and Vere-Jones, 2003)). Therefore, a prediction of the model is a strong link between differences in dynamical properties of states and differences in membrane potential statistics, with the latter changing systematically from non-Gaussian to Gaussian across different regimes. This was validated in experimental data (we observed a linear decrease of *γ_V_* with *μ_V_*; **Figure 5C**).

Finally, our model makes predictions about how a cortical circuit may respond to a strong sustained afferent input (**Figure 3C**) that emulates the input driving primary sensory cortices upon the static presentation of an optimal receptive-field stimulus. Recent experimental observations in the awake mammalian cortex showed three prominent phenomena in response to this type of sensory stimuli. First, in the mouse visual cortex, the presentation of a grating was associated with the predominance of the recurrent component compared to the afferent (thalamic) component of synaptic excitation (Reinhold et al., 2015), together with a fast time scale for the relaxation of the network activity (*τ_NTWK_* ~ 10 ms). Second, such a stimulus also led to the transient prevalence of inhibitory conductances (Haider et al., 2013), a situation consistent with balanced currents given the asymmetry of the driving forces for excitatory and inhibitory currents. Third, in the monkey visual cortex the presentation of a grating triggered a transition to Gaussian membrane potential fluctuations (Tan et al., 2014). In light of the model presented in this study, those experimental observations are natural correlates of the stimulus-evoked dynamics: a strong afferent excitatory input transiently switches network activity toward a network regime where recurrent input dominates synaptic activity (**Figure 1C**), the relaxation time constant is low (**Figure 3F**), depolarized fluctuations become Gaussian (**Figure 3G**), and inhibitory conductances dominate (**Figure 1K**).

### Hypothetical functions of the various non-rhythmic waking states suggested by the model

The transition toward aroused or attentive states elicits desynchronization of network activity in sensory cortices (Harris and Thiele, 2011). This mechanism is thought to facilitate sensory processing as sensory-evoked activity exhibits higher signal-to-noise ratio when low-frequency cortical rhythm fades out (Busse et al., 2017; Harris and Thiele, 2011). However, the functional modulation of sensation beyond the “desynchronization” effect, i.e. within the various non-rhythmic substates of wakefulness, remains elusive. Our computational model provides new mechanistic insight on this process. It suggests that neocortical networks switch their encoding properties upon modulation of the afferent excitatory input in order to either faithfully encode complex patterns of presynaptic activity (in the AD regime) or to exhibit strong population-wide recurrent amplification of any afferent input (in the RD regime). Experimental results in the mouse auditory and visual cortices seem to match this model prediction. In fact, the behavioral state of the animal, indexed based on pupil size and running speed into low arousal, moderate arousal, and hyper arousal, was shown to modulate the *V_m_* signature of cortical dynamics similarly to what observed in the model ((McGinley et al., 2015b; Reimer et al., 2014), see (Busse et al., 2017) for a review). Comparing the experimental data presented in those studies with the predictions of our study, the AD regime could correspond to the moderate arousal state (characterized by low depolarization level, non-rhythmic membrane potential dynamics associated to intermediate pupil size) while the RD regime could correspond to the hyper arousal regime (characterized by depolarized fluctuations of the membrane potential of narrow variance associated to locomotion and/or large pupil size). Interestingly, moderate arousal was found to be optimal for the discrimination of a tone-in-noise auditory stimulus (McGinley et al., 2015b), a result in accordance with the model prediction of a more reliable subnetworks activation (tone-specific in this context) in the AD regime (**Figure 6D**). A theoretical explanation for the inverted U-relationship for detection performance with a peak at moderate arousal (McGinley et al., 2015a, 2015b) might thus be found in the reliable encoding of sensory stimuli characterizing the AD regime. During locomotion (hyper arousal), neuronal responses in the visual system were found to be increased at all orientations (Reimer et al., 2014), consistent with the prediction of an unspecific recurrent amplification of population activity in the RD regime (**Figure 6H**). Neuronal responses of preferred orientation were selectively potentiated only in the moderate arousal state (Reimer et al., 2014).

Because the precise spatiotemporal pattern of neural responses within sensory cortices is thought to encode stimulus identity (Luczak et al., 2015; Panzeri et al., 2010), the AD regime might be an activity regime optimized for sensory discrimination. In contrast, the fast and unstructured amplification of excitatory inputs that characterizes the RD regime may potentiate the cortical response to weak sensory stimuli and could therefore represent an activity regime optimized for sensory detection. Because the electrophysiological signature of the RD regime is found under hyper arousal conditions (McGinley et al., 2015a) and it is presumably associated to a high cognitive load, this state might enable animals to focus on a narrow piece of information, e.g. the presence (or absence) of a sensory cue within the environment. Future work focusing on the modulation of sensation in awake behaving animals across various sensory modalities will test the validity and generality of this theoretical framework.

## Material and Methods

### Animals

Experimental procedures involving animals have been approved by the IIT Animal Welfare Body and by the Italian Ministry of Health (authorization # 34/2015-PR and 125/2012-B), in accordance with the National legislation (D.Lgs. 26/2014) and the European legislation (European Directive 2010/63/EU). Experiments were performed on young-adult (4-6 weeks old, either sex) C57BL/6J mice (Charles River, Calco, Italy) and PV-IRES-Cre mice (B6.129P2-^Pvalbtm1(cre)Arbr/J^, Jackson Laboratory, Bar Harbor, USA). The animals were housed in a 12:12 hr light-dark cycle in singularly ventilated cages, with access to food and water *ad libitum*.

### In vivo electrophysiology in awake mice

The experimental procedures for *in vivo* electrophysiological recordings in awake head-fixed mice have been previously described (Zucca et al., 2017). Briefly, a custom metal plate was fixed on the skull of young (P22-P24) mice two weeks before the experimental sessions. After a 2-3 days recovery period, mice were habituated to sit quietly on the experimental setup for at least 7-10 days (one session per day and gradually increasing session duration). The day of the experiment, mice were anesthetized with 2.5% isofluorane and a small craniotomy (0.5 mm × 0.5 mm) was opened over the somatosensory cortex and a 30 minutes long recovery period was provided to the animal before starting the recordings. Brain surface was kept moist with a HEPES-buffered artificial cerebrospinal fluid (ACSF). Current-clamp patch-clamp recordings were carried out on superficial pyramidal neurons (100 – 350 μm). 3–6 MΩ borosilicate glass pipettes (Hilgenberg, Malsfeld, Germany) were filled with an internal solution containing (in mM): K-gluconate 140, MgCl_2_ 1, NaCl 8, Na_2_ATP 2, Na_3_GTP 0.5, HEPES 10, Tris-phosphocreatine 10 to pH 7.2 with KOH. For simultaneous recordings of multi-unit activity (**Figure S5**), an additional glass pipette filled with ACSF was lowered into the tissue with the deeper tip placed at ^~^300 μm from pial surface. Electrical signals were acquired using a Multiclamp 700B amplifier, filtered at 10 kHz, digitized at 50 kHz with a Digidata 1440 and stored with pClamp 10 (Axon Instruments, Union City, CA). We recorded from n = 14 cells in N = 4 Wild Type (WT) C57BL/6J animals. In the analysis, we added data from n = 8 cells in N = 4 PV-Cre mice obtained in recordings that were designed for a previous publication (Zucca et al., 2017). Those recordings contained period of optogenetic stimulation (every 5 s, see details in Zucca et al., 2017) of PV cells intermingled with period of spontaneous activity. The stimulation epochs and subsequent 500 ms-long time periods were discarded from the analysis in the additional 8 cells of PV-Cre mice. All the relations displayed in Figure 4 for the pooled data (WT + PV-Cre) were found similarly significant in the dataset containing only the WT mice (p < 1e-3 for all relations with similar correlation coefficients, see **Figure S4C**).

### Computing the electrophysiological properties of non-rhythmic epochs

From the previously described recordings, we extracted stable membrane potential samples. Cells or periods with action potential peaking below 0mV or displaying a slow (~1min) drift in the *V_m_* trace were discarded from the analysis. This resulted in dataset of n=22 cells with a recording time per cell of 5.1±3.2 min. This stability criterion enabled us to perform the analysis on an absolute scale of membrane potential values (see **Figure 4** and **Figure S4**).

We first estimated a time-varying low frequency power within the *V_m_* samples: 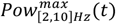. To this purpose, we discretized the time axis over windows of 500ms sliding with 25ms shifts and extracted the maximum power within the [2,10]Hz band (estimated with a fast Fourier transform algorithm, <u>numpy.fft</u>). All segments whose center *t_i_* had a 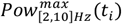 value greater than the *rhythmicity threshold* were classified as “rhythmic” and discarded from future analysis. The value of the *rhythmicity threshold* was adjusted so that 50% of the data should be classified as “rhythmic” (see **Figure 4C**, in **Figure S4D** we analyze various *rhythmicity threshold* levels). In the remaining “non-rhythmic” samples {*t_i_*}_*NR*_, we evaluate the mean depolarization level *μ_V_* (*t_i_*) over the same 500ms interval surrounding the center time *t_i_* (T=500ms is a good tradeoff between an interval short enough to catch the potential variability in network regimes at the sub-second time scale, i.e. T<1s, and an interval long enough to overcome the relaxation time of the network dynamics, i.e. T ≫10ms, see main text). At that point, each time *t_i_* is associated to a given depolarization level *μ_V_* (*t_i_*). We now discretize the *μ_V_* axis in *j* ∈[1,20] points from −80mV to −50mV and we count the number of segments *n_j_* over all *t_i_* where 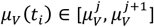 (see **Figure S4A**). As all cells did not contribute equally to all *μ_V_* levels (see **Figure S4C**), we applied a “minimum contribution” criteria: if a depolarization level counted less than 200 segments (*n_j_*<200), the 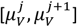 level was discarded from future analysis (see **Figure S4A**). We then count the number of spikes falling in a given level 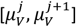 level by counting spikes within the 500ms window. Spikes were detected as a positive crossing of the −30mV level (spikes were blanked in the *V_m_* traces by discarding the values above this threshold). We then compute the fluctuations properties of all depolarization levels. This was achieved by constructing a “pooled distribution” and a “pooled autocorrelation function” corresponding to all 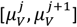 intervals. For all 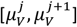 intervals, we took 500ms samples around all *t_i_* matching 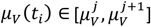 and incremented the “pooled distribution” with those *V_m_* samples. Similarly, we incremented the “pooled autocorrelation function” with the individual normalized autocorrelation functions (evaluated up to 100ms time shift) of those *V_m_* samples. The resulting “pooled distributions” and “pooled autocorrelation functions” are illustrated for a single cell on **Figure S4A**. The “pooled distributions” at all 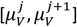 levels were used to evaluate the standard deviation 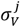 and skewness 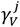 while the “pooled autocorrelation functions” were used to determine the *autocorrelation time* 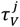. The *autocorrelation time* 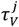 was determined by a numerical integration of this normalized autocorrelation function (Zerlaut et al., 2016). This procedure was repeated for all cells (shown in **Figure S4B**) and yielded the population data of **Figure 4**. We also analyzed the goodness-to-fit of a Gaussian fitting of the “pooled distributions”, we performed a least-square fitting (using the function scipy.optimize.leastsq) and we report the coefficient of determination *R*^2^ (see main text).

### Numerical simulations of recurrent network dynamics

We studied two versions of recurrently connected networks targeted by an afferent excitatory population: 1) a model with two coupled populations (excitatory and inhibitory neurons) and 2) a three population model with excitatory, inhibitory and disinhibitory neurons. Single cells were described as single compartment Integrate and Fire models with conductance-based exponential synapses. Their membrane potential dynamics thus follows the set of equations:

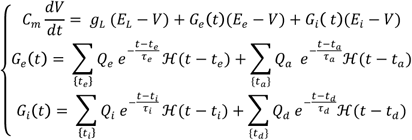

Where ℋ is the Heaviside (step) function. Note that, to emphasize the similarity in the equation between the different cell types considered (excitatory, inhibitory and disinhibitory), we omitted the index of the target cell (e.g. the weight should be *Q_ae_* for the afferent excitation onto the excitatory cell instead of *Q_a_* here). This set of equation is complemented with a threshold and reset mechanism, i.e. when the membrane potential *V* reaches a threshold *V_thre_* it is reset at the value *V_reset_* during a refractory period *τ_refrac_*. The sets of events {*t_X_*}corresponds to the synaptic events targeting a specific neuron. All parameters can be found on **Table 1** for the two population model (**Figure 1**). The additional parameters required for the coupled three population model (excitation, inhibition, disinhibition, for **Figure 3** and **Figure 6**) can be found on **Table 2**.

Recurrent connections were drawn randomly by connecting each neuron of the population *Y* with *p_XY_N_X_* neurons of the population *X*. Afferent drive of frequency *v_a_* onto population *X* with connectivity probability *p_aX_* was modeled by stimulating each neuron of the population *X* with a Poisson process of frequency *p_aX_N_a_v_a_* (i.e. using the properties of Poisson processes under the hypothesis of independent processes).

Numerical simulations were performed with the <u>Brian2</u> simulator (Goodman and Brette, 2009) (RRID:SCR_002998). A time step of dt=0.1ms was chosen. Stationary properties of network activity were evaluated with simulations lasting 10s. The first 200ms were discarded from the analysis to remove the contributions of initial transients. Simulations were repeated over multiple seeds generating different realizations of the random connectivity scheme and of the random afferent stimulation (see number in the legends).

### Characterizing network dynamics

From the numerical simulations, we monitored all spike times and binned them in *T_b_*=2ms time bins to obtain the spike train *S_i_*(*t*) for each neuron *I* (*S_i_*(*t*) takes only 0 or 1 values as *τ_refrac_* > *T_b_*) We analyzed the network activity by looking at the time-varying firing rate of the population *X*:

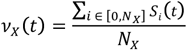

We measured population synchrony by averaging the correlation coefficient of the spike trains over some (*i,j*) neuronal pairs (Kumar et al., 2008), i.e. the synchrony index *SI* was given by:

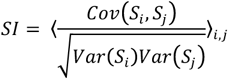

In practice we selected 4000 spiking neuronal pairs for numerical evaluation.

Additionally, we monitored the membrane potential, the synaptic conductances and the synaptic currents of four randomly chosen cells in each populations. To evaluate the mean, standard deviations, skewness and autocorrelation time of the membrane potential fluctuations, we discarded the refractory periods from the analysis. The same discarding procedure was applied for the mean conductances and currents reported here. The excitatory currents and conductances shown in the main text merge all excitatory contributions together (afferent and recurrent excitations). The inhibitory currents and conductances correspond to recurrent inhibition only for excitatory cells and add the disinhibitory contributions for inhibitory cells in the three population model.

### Varying parameters of the network model

We investigated the robustness of the proposed theoretical picture by studying its sensitivity to parameter variations. The values of parameters and results of this analysis is shown on **Figure S3**. Network simulations were run with time step 0.1ms, lasted 10s and were repeated over 4 different seeds.

### Response to an afferent time-varying rate envelope

To emulate a time-varying afferent input onto the local cortical network (see **Figure 3**), we took an arbitrary waveform for the firing rate activity of the afferent population. From this waveform, an inhomogeneous Poisson process was generated to stimulate each neuron of the three populations model. For **Figure 3**, the waveform was taken as:

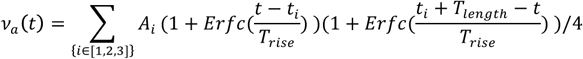

with *A*_1_=4Hz, *A*_2_=18Hz, *A*_3_=8Hz, *t*_1_=100ms, *t*_2_=1150ms, *t*_3_=2000ms, *T_rise_*=50ms and *T_length_*=900ms. The resulting waveform is shown in **Figure 3C**.

### Determining the relaxation time constant of the network dynamics

We determined the network time constant *τ_NTWK_* at three different levels of network activity in the three-population model (see **Figure 3F**). The network model was stimulated with three different levels of stationary background activity *v_a_* = 4 Hz, *v_a_* = 8 Hz and *v_a_* = 18 Hz. On top of this background activity, we added a 2Hz step of afferent excitation lasting *T_stim_*=100ms and each 500ms. We repeated this stimulation a 100 times and we computed the trial-average response to this stimulus (shown in the inset of **Figure 3F**). The network time constant was estimated by a least-square fitting of the following waveform: 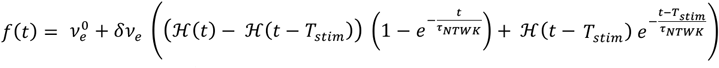. The three values 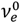, *δv_e_* and *τ_NTWK_* were determined through the minimization procedure. We show the *τ_NTWK_* values in the bar plot and the response amplitudes *δv_e_* as the scale bar annotations in **Figure 3F**.

### Encoding of spiking patterns of presynaptic activity

We designed a stimulation to investigate whether a complex spatiotemporal pattern targeting a subset of the local cortical population could be faithfully encoded by the activity of this sub-network (see **Figure 6A-E**). We took the following scheme. Within the 100 neurons of the afferent population, we made groups of 10 neurons that co-activate simultaneously. Those groups of 10 neurons target a subset of 100 neurons within the 4000 neurons of the excitatory population (with a synaptic weight equal to those of background afferent connections). Presynaptic neurons only make mono-synaptic connections to a target neuron, but two co-activated neurons may connect to the same neuron in the 100 neurons target population hence creating some degree of synchronous activation (but with a low probability: 1%, because *p_ae_*=10%, see **Table 1**). Within a window of 500ms, we generate random activations over time with a homogeneous Poisson process of frequency 20 Hz (i.e. 10 activations per 500 ms window) and assign randomly each activation time to a given afferent group, this generates one pattern (see example patterns on **Figure 6A** and **Figure S6A**). We reproduce this procedure 10 times with a different random seed to obtain 10 patterns of presynaptic activations. We then feed the network with this afferent pattern on top of the non-specific background afferent drive (both in the AD regime and in the DR regime). We run 20 trials per pattern, where the realization of the background activity varies while the pattern is kept constant. We compared the output spiking patterns using the inner-product (*IR*) and distance (*D*) for multi-neuron spike trains derived in (Houghton and Kreuz, 2012; Rossum, 2001) implemented in the publicly-available package <u>pymuvr</u>. This metrics takes a time scale *τ* that sets the temporal sensitivity (for *τ* → 0 the metrics is only sensitive to infinitely precise coincident spiking, for *τ* → ∞ the metrics is a joint spike count over time). The value of *τ* was set to 5 ms as this time scale was found in preliminary analyses to be the minimal time scale for which a reliable encoding of the input pattern was observed in the AD regime. The cross-correlation coefficient between spike trains computed in **Figure 6D** was computed as 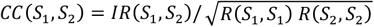 where *IR* is the inner product between two spike trains *S*_1_ and *S*_2_. We then implemented a k-neighbor classifier to decode the output spike train of the excitatory subnetwork. The distance between two output patterns relied on the distance *D*. We implemented this custom metrics in the k-neighbor-classifier of <u>scikitlearn</u> (Pedregosa et al., 2011)(RRID:SCR_002577) to obtain our classifier. We first train the classifier on the first 10 trials and tested on the last 10 trials per pattern. For a first-nearest-neighbor classification, we found the following decoding accuracies: 88.0 ± 9.8 % for AD activity and 16.0 ± 0.2 % for RD activity. Raising the number of neighbors up to 10 points (over a training set containing 10 trials per presynaptic patterns) did not affect this difference: it yielded (non-monotonic) variations of the decoding accuracy between 90% and 65% for AD regimes and between 27% and 16% for RD regimes, we therefore kept a nearest-neighbor classifier for all analysis. To partially separate the spatial and temporal components in afferent patterns encoded by the network activity, we duplicated all 10 patterns and their 10 repetitions in the training set by aligning the network response onset time to all observed stimulus onsets (shifting the time axis, the procedure is depicted in **Figure S6A**). The final decoder should therefore associate a trial in the test set with a given pattern identity and a given stimulus onset (accuracy results shown in **Figure 6E**). In **Figure S6B**, we show the distributions of decoded stimulus onsets in the AD and RD regimes. The percentage of stimulus-evoked activity (see main text) was evaluated by comparing the firing rates in the 500 ms before and during the 500 ms of the stimulus.

### Encoding of presynaptic rate waveforms

We designed a stimulation to investigate whether the rate envelope of given presynaptic stimulus could be faithfully encoded by the activity of the network (see **Figure 6F-J**). The waveform was taken as:

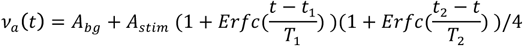

with *T*_1_=100ms, *T*_2_=300ms, *t*_1_=400ms and *t*_2_=1100ms. *A_bg_*=4Hz to produce the AD regime and *A_bg_*=14Hz to produce the DR regime. *A_stim_* was varied from 0.1Hz to 7Hz in 5 different levels (see the resulting waveforms are shown in **Figure 6F**). This time-varying rate was then converted to a Poisson process (varying the seed in all trials) setting the activity of the afferent population and fed as an input to the recurrent network. Similarly to the previous section, we implemented a k-neighbor classifier to decode the rate waveform from a sub-population of the network (taking the same 100 neurons sample). The time-varying rate of the subpopulation was computed by binning spikes in 2ms bins and Gaussian smoothing of extent 30ms, yielding the quantity *R*(*t*). The metric for the rate waveform decoder was the integral over the stimulus duration of the square difference between waveforms, i.e. for two waveforms R1 and R2, it corresponded to:

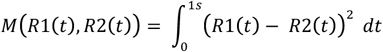

We run 20 trials for each of the five levels of afferent activity shown in **Figure 6F**. We trained the decoder on the first 10 trials and tested it on the following 10 trials.

### Data Analysis and Statistics

Data were analyzed with <u>SciPy</u> (Oliphant, 2007). Experimental data were translated to the Python format using Neo (Garcia et al., 2014) (RRID:SCR_000634). In **Figure 5A-D**, we performed least-square linear regressions on continuously distributed data (implemented in <u>scipy.stats.linregress</u>), we report the correlation coefficients (“c”). Given the partial temporal overlap between individual samples of membrane potential, the data across the different *μ_V_* levels cannot be considered as independent, so we evaluated statistical significance (“p”) with a non-parametric one-tailed permutation test (performed with 1e5 permutations, hence p values were reported as “p<1e-5” if no permutation was found to exhibit the correlation value of the data). In **Figure 6**, we evaluated the significance of the difference in encoding accuracy and response gain with a two-sided t-test (implemented in <u>scipy.stats.ttest_rel</u>).

## Acknowledgments

We wish to thank Eugenio Piasini for advice on the decoding analysis and Diego Fasoli for comments on the manuscript. Research funded by the European Research Council (NEURO-PATTERNS), the FP7-602531 (DESIRE), and, in part, the Flag-Era Joint Transnational Call (SLOW-DYN).

## Supplementary Information

The derivation of the *mean-field* analysis is presented on the **Supplementary Information** with four supplementary figures. The code for the numerical simulations and analysis producing the main and supplementary data is publicly available in the form of an *Interactive Python notebook* (Pérez and Granger, 2007) on the following link: https://github.com/yzerlaut/notebook_papers/blob/master/The_Spectrum_of_Asynch_Dynamics_2018.ipynb.

## Authors Contribution

Conceptualization, Y.Z., S.Z., S.P., T.F.; Investigation, Y.Z., S.Z.; Formal Analysis and Software, Y.Z.; Writing, Y.Z., S.Z., S.P., T.F.; Funding Acquisition, S.P., T.F.; Supervision, S.P., T.F.

